# Concurrent *PIK3CA* mutant drives cachexia through inflammatory signaling in *EGFR* mutant lung cancer

**DOI:** 10.1101/2025.01.17.633515

**Authors:** Meiting Yue, Zhen Qin, Shijie Tang, Xinlei Cai, Yikai Zhao, Liang Chen, Luonan Chen, Hongbin Ji

## Abstract

*PIK3CA* mutation is frequently concurrent with known oncogenic drivers such as *EGFR* mutation in lung cancer, raising an interesting question about its real function. Cachexia is a systemic disease involving complex interaction between primary tumors and distant organs, significantly contributing to cancer-related mortality. Through integrative study of genetically engineered mouse models (GEMMs) and clinical data, we find concurrent *PIK3CA* mutant preferentially drives cachexia in *EGFR*-mutant lung cancer, promoting malignant progression instead of cancer initiation. *PIK3CA* mutant-mediated cachexia could be overcome by osimertinib (Osi) treatment in Osi-sensitive GEMM. In contrast, chemotherapy, routinely used in clinic for those relapsed from Osi therapy, fails to ameliorate cachexia in Osi-resistant GEMM despite notable tumor suppression. *PIK3CA* mutant-driven cachexia is mediated through NF-κB activation and could be dampened by combined aspirin treatment. This work uncovers the biological function of *PIK3CA* mutant and mechanism behinds its clinical impacts, and proposes a potentially effective strategy for clinical management.

## Introduction

*PIK3CA*, encoding the p110α catalytic subunit of PI3K, is mutated in approximately 7-16% of non-small-cell lung cancer (NSCLC)^1,2^. *PIK3CA* mutations are highly enriched in its helical region (*E542K*, *E545K*) and kinase domain (*H1047R*)^3^. It’s well established that these mutations are oncogenic and frequently cause aberrant activation of downstream AKT signaling, triggering a cascade of responses that drive lung tumorigenesis^4^. Animal model studies show that *PIK3CA* mutant transgene alone is sufficient to drive lung cancer initiation in mice^5^. Paradoxically, most *PIK3CA* mutations are found to occur concurrently with those famous oncogenic drivers^6,7^, e.g., *EGFR* mutations, *KRAS* mutations and *ALK* fusions, which are known to be mutually exclusive^8-11^ and individually suffice to drive lung cancer initiation^12-14^.

Previous study has shown that approximately 80% of lung cancer harboring *PIK3CA* mutations tends to have *EGFR* mutations in East Asian cohort^7^. Osimertinib (Osi), a third-generation EGFR tyrosine kinase inhibitor (TKI), is currently the preferable option for *EGFR*-mutant NSCLC patients, owing to its high efficacy and well-tolerated safety profile^15^. However, similar to that of early-generation EGFR-TKIs, Osi resistance inevitably develops. Additional *C797S* mutation represents one of the most prevalent mechanisms in Osi resistance^16^. When occurring in conjunction with a sensitizing mutation and in the absence of *T790M* mutation, *C797S* mutation confers resistance to Osi while preserving sensitivity to first- and second-generation agents. The presence of triple mutants consisting of the sensitizing mutation, *T790M*, and *C797S*, leads to resistance against all three generations of EGFR TKIs^17^. For these patients, chemotherapy, e.g., pemetrexed (PEM) in combination with cisplatin (CDDP), is the remaining option in clinic^18^.

High-throughput sequencing analyses of TKI-resistant lung cancer specimens have identified the emergence of additional *PIK3CA* mutation^19,20^, indicative of its potential contribution to drug resistance. This is further supported by the observation of worse overall survival (OS) of those *EGFR*-mutant patients with concurrent *PIK3CA* mutation in their tumors^21,22^. However, detailed analyses of clinical TKI responses show that the concurrence of *PIK3CA* mutation does not affect the therapeutic efficacy of TKI treatments^21,22^, e.g., almost no difference of progression-free survival (PFS) in patients with or without *PIK3CA* mutations. This raises another interesting question regarding the real mechanism behinds the clinical impacts of *PIK3CA* mutant.

Cancer cachexia is a complex and debilitating syndrome characterized by body weight loss primarily due to skeletal muscle atrophy and adipose tissue wasting^23^, responsible for more than 20% of cancer-related deaths^24^. Cachexia occurs in approximately 50% of lung cancer patients^25^ and is associated with poor life quality^26^, elevated treatment-related toxicity^27,28^, reduced therapeutic responses^29^ and increased risk of mortality^30-33^. Recent study has begun to focus on the link between specific gene alterations and cachexia development^30^. Through the integrative analyses of genetically engineered mouse models (GEMMs) and human clinical data, we here uncover an unexpected biological function of *PIK3CA* mutations in lung tumorigenesis, mainly contributing to cachexia instead of driving lung cancer initiation. Moreover, we also provide a reasonable explanation about the clinical observation of paradoxical difference between PFS and OS in link to concurrent *PIK3CA* mutation, and propose a potentially effective strategy for clinical management of lung cancer patients with concurrent *PIK3CA* mutation.

## Results

### *PIK3CA* mutant contributes to malignant progression but not lung cancer initiation

To investigate the real function of *PIK3CA* mutant in lung tumorigenesis, we generated a genetically engineered mouse model (GEMM) by integrating the LoxP-stop-LoxP-*PIK3CA E545K* transgene into the Rosa26 locus of C57BL/6 mice (Fig. S1A), in which the *PIK3CA* mutant can be conditionally expressed via intranasal inhalation of Ad-Cre virus^34^. Unexpectedly, we found no tumor formation even after 40 weeks of Ad-Cre administration (Fig. S1B-C). This indicates that *PIK3CA* mutant expression alone seems insufficient to drive lung cancer initiation. Given that *PIK3CA* mutation and *TP53* mutation frequently co-occur in human NSCLC (Fig. S1D), we further crossed *PIK3CA^E545K^* to *Trp53^flox/flox^* mouse to see if there is any lung tumor formation. Again, we found no tumor formation in this model (Fig. S1E-F). These data collectively point to a dispensable role of *PIK3CA* mutant in driving lung cancer initiation.

It’s well established that either *EGFR* mutation*, KRAS* mutation, or *ALK* fusion alone suffice in driving lung cancer initiation in mice^12-14^. *EGFR* mutation is the oncogenic driver most significantly associated with the concurrence of *PIK3CA* mutation (Fig. S2A). We then crossed *PIK3CA^E545K^*mice with *EGFR^L858R^;Trp53^flox/flox^* (EP) mice to generate the *EGFR^L858R^;PIK3CA^E545K^;Trp53^flox/flox^* (EPP) cohort for further study (Fig. S2B). Both *in vitro* and *in vivo* assays demonstrate that *PIK3CA* mutant can activate the downstream AKT signaling and promotes lung cancer progression (Fig. S2C-K). Further analyses of *in situ* tumors showed that *PIK3CA* mutant significantly promoted lung cancer malignant progression, characterized by a greater degree of cellular pleomorphism and nuclear atypia^35^ (Fig. S2L-M). The tumor-bearing EPP mice exhibited significantly reduced survival compared to the EP mice (Fig. S2N). These data collectively demonstrate that *PIK3CA* mutant mainly contributes to lung cancer malignant progression instead of driving lung cancer initiation.

### *PIK3CA* mutant drives cachexia in *EGFR* mutant lung cancer

Interestingly, the tumor-bearing EPP mice displayed a rapid decrease in body weight starting from 6 weeks post Ad-Cre treatment, which was not observed in either wild type (WT) or EP control mice (Fig. 1A-B). Further analyses of the body composition revealed that the EPP mice experienced a continuous loss of lean mass along with a decrease in fat mass, aligning with the clinical diagnostic criteria for cachexia as previously established^36^ (Fig. S3A-B). Moreover, fat imaging visually illustrated a significant decrease of fat tissue in these tumor-bearing EPP mice (Fig 1C). Compared to WT and EP mice, the EPP mice exhibited significant decrease in muscle mass in gastrocnemius (GAST), tibialis anterior (TA), and quadriceps (QUAD), as well as a decrease in epididymal white adipose tissue (eWAT) in males and the gonadal white adipose tissue (gWAT) in females (Fig. 1D-E). H&E staining analyses revealed the reduction of muscle fiber cross-sectional area (CSA) and adipocyte size in the EPP mice (Fig. S3C-G), histologically confirming these atrophic changes.

**Fig. 1.**
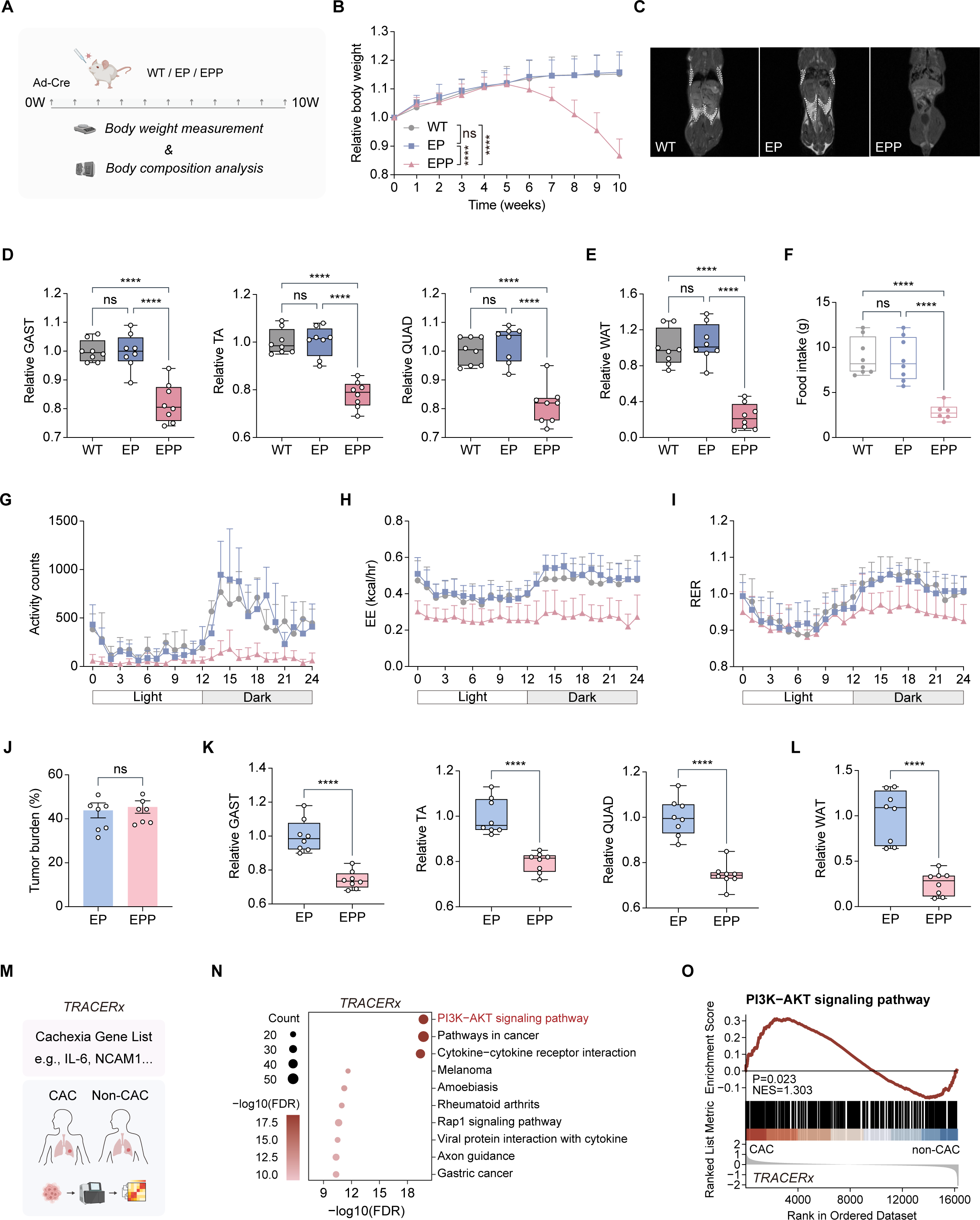
P*I*K3CA mutant is associated with cachexia development in lung cancer. A. Schematic diagram of long-term monitoring strategy of mouse body weight and body composition. B. Relative changes in body weight of WT mice (n=12), EP mice (n=12), and EPP mice (n=12) over time following intranasal delivery of Ad-Cre for 0-10 weeks. Data were normalized based on the values at the time of intranasal induction, and data collected at the 10-week post-induction were utilized for differential analysis. C. Representative fat imaging of WT mice, EP mice, and EPP mice following 8 weeks post Ad-Cre intranasal delivery. D-E. Assessed weights of the gastrocnemius (GAST), tibialis anterior (TA), quadriceps (QUAD) (D), and white adipose tissue (WAT) (E) in WT mice (n=8), EP mice (n=8), and EPP mice (n=8) following 8 weeks post Ad-Cre intranasal delivery. Data were normalized based on the average values of corresponding tissue in WT mice with the same gender. F. Average daily food intake of WT mice (n=8), EP mice (n=8), and EPP mice (n=6) following 8 weeks post Ad-Cre intranasal delivery. G-I. 24-hour activity counts (G), energy expenditure (EE) (H), and respiratory exchange ratio (RER) (I) of WT mice (n=8), EP mice (n=8), and EPP mice (n=6) following 8 weeks post Ad-Cre intranasal delivery. J. Statistical analyses of tumor burden in EP mice (n=8) and EPP mice (n=8). K-L. Assessed weights of the gastrocnemius (GAST), tibialis anterior (TA), quadriceps (QUAD) (K), and white adipose tissue (WAT) (L) in EP mice (n=8) and EPP mice (n=8). Data were normalized based on the average values of corresponding tissue in EP mice with the same gender. M. Schematic diagram of bioinformatics analysis base on TRACERx study (Al-Sawaf *et al*., 2023). N. Enrichment of cachexia candidate gene list in KEGG pathways (TRACERx study, Al-Sawaf *et al*., 2023). The size of the dots reflects the number of enriched genes in the pathway. The color of the dots indicates the significance of the enrichment. O. GSEA enrichment plot of PI3K-AKT signaling pathway in cancer-associated cachexia (CAC) patients compared to non-CAC patients in TRACERx study (Al-Sawaf *et al*., 2023) cohort. Data are presented as mean ± SD. *P < 0.05, ***P < 0.001,****P< 0.0001 by one-way ANOVA (B, D-F), two-tailed unpaired Student’s t test (J-L). ns: not significant.

Metabolic monitoring revealed that the EPP mice exhibited a significant decrease of food intake, lower level of locomotor activity, and reduced energy expenditure (EE) (Fig. 1F-H), all of which are hallmarks of cachexia in clinical settings^25^. Moreover, the lower respiratory exchange ratio (RER) in EPP mice indicated a metabolic shift toward the utilization of non-glucose substrates, potentially related to their decreased food intake (Fig. 1I). These characteristics were particularly pronounced at night, coinciding with the activity rhythms of mice (Fig. S3H-J).

To exclude the potential impact of tumor burdens upon cachexia, we further analyzed the EP mice and EPP mice with comparable tumor burdens at different time points post Ad-Cre treatment, e.g., the EP mice post 17 weeks of Ad-Cre treatment and the EPP mice post 10 weeks of Ad-Cre treatment (Fig. 1J). Interestingly, a significant difference of cachexia features was still detectable in these two groups, e.g., the EPP mice showed lower levels of mass of skeletal muscles and adipose tissues (Fig. 1K-L). These data demonstrate that *PIK3CA* mutant triggers the cachexia symptom independent of tumor burdens.

Analyses of cachexia signature proposed by *TRACERx* study^30^ highlighted the PI3K-AKT signaling as one of the most significantly enriched pathways (Fig. 1M-N). Compared to patients without cachexia, the PI3K-AKT signature was also significantly enriched in those primary tumors from patients with cachexia (Fig. 1O). Consistently, we observed that PI3K-AKT signaling was similarly enriched in cachectic individuals across two additional datasets of lung tumor RNA sequencing (RNA-seq)^37^ and serum proteomics^38^ (Fig. S4A-B). These data collectively support the significant role of *PIK3CA* mutant in driving cachexia.

### Muscle wasting characteristics in EPP mice recapitulate clinical phenomena

Muscle wasting is the most critical characteristic of cancer cachexia^36^. We next comparatively analyzed the gene expression profiling of TA muscles derived from WT, EP, and EPP mice (Fig. 2A). Principal component analysis (PCA) revealed separation between TA muscles from EPP mice and those from WT and EP mice (Fig. 2B). Consistently, gene expression pattern of EPP mice differed dramatically from the others (Fig. 2C). We found an enrichment of proteasome and autophagy signaling in TA muscles of EPP mice (Fig. 2D), which has been extensively documented in muscle atrophy^39^. An enrichment of features associated with muscle disorder-related diseases was also observed, suggesting that muscle atrophy in cancer cachexia may share signaling pathways with other muscle disorders.

**Fig. 2.**
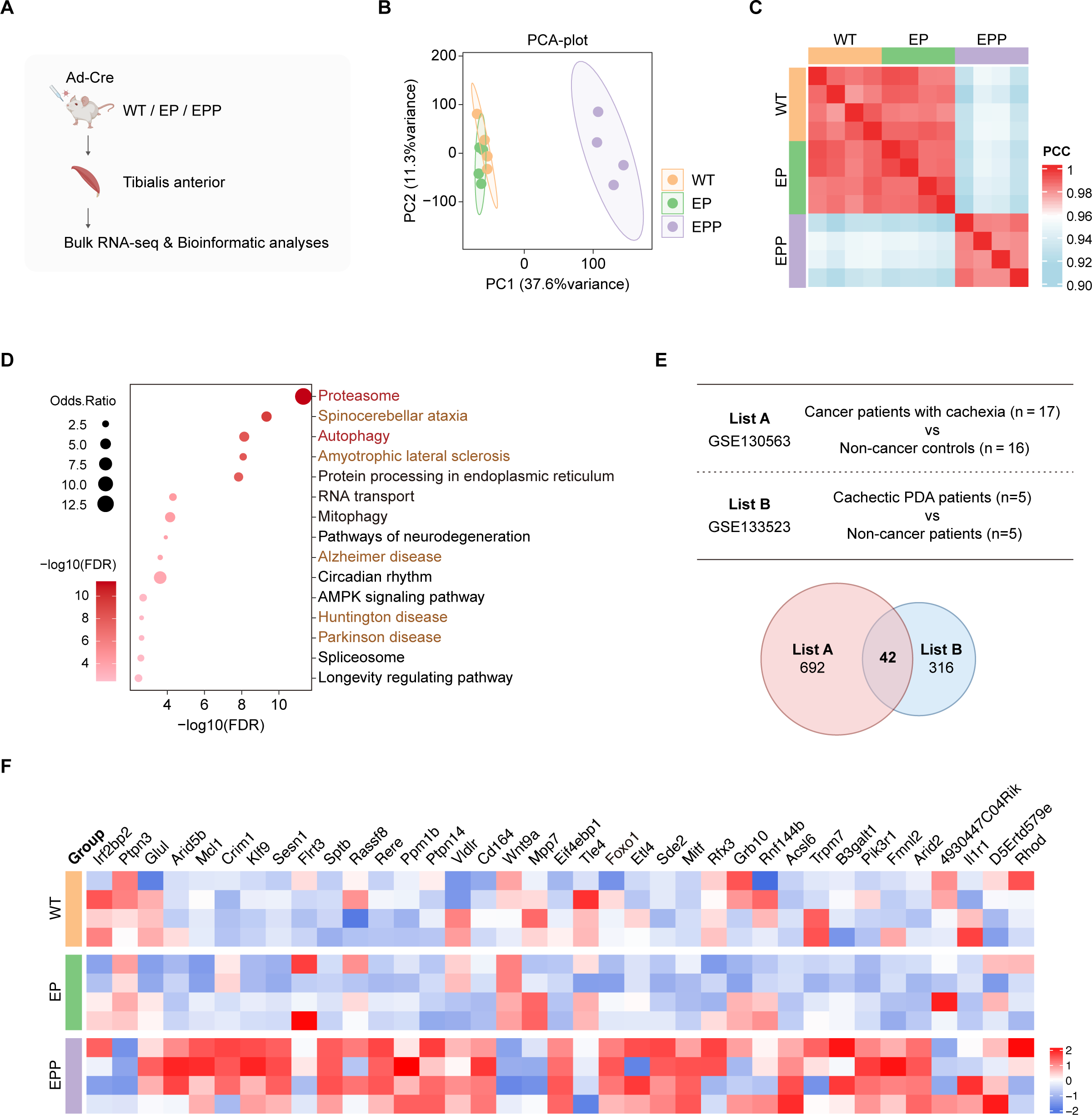
Characteristics of muscle wasting in cachectic mice recapitulate those observed in patients with cachexia. A. Schematic diagram of murine skeletal muscle RNA sequencing. B-C. Principal component analysis (PCA) plot of tibialis anterior tissues from WT mice (n=4), EP mice (n=4), and EPP mice (n=4). C. Heatmap showing Pearson correlation coefficient (PCC) among tibialis anterior tissues from WT mice (n=4), EP mice (n=4), and EPP mice (n=4). D. Dot plot shows enriched KEGG pathways within the tibialis anterior tissues of EPP mice compared to EP mice, based on significantly differentially expressed genes (Cutoff: FDR < 0.05 and fold change ≥ 1.5). The size of the dots reflects the odds ratio of enriched genes in the pathway. The color of the dots indicates the significance of the enrichment. E. Left part contains database information; right part displays a Venn diagram showing the number of upregulated genes in muscle tissue of patients with cancer cachexia compared to non-cancer controls. F. Gene expression heatmap of tibialis anterior tissues from WT mice (n=4), EP mice (n=4), and EPP mice (n=4).

We next analyzed two public databases (GSE130563^40^; GSE133523^41^) containing expression data from muscle tissues of cachectic patients and their respective controls. We found 42 genes upregulated in the muscles of cachectic patients, with 37 genes shared homologous counterparts in murine tissues with detectable expression (Fig. 2E). We further found that most of these genes were enriched in the muscle tissues of EPP mice (Fig. 2F), suggesting that the muscle wasting features in these mice closely resemble clinical observations.

### Osi effectively inhibits lung cancer progression and alleviates cachexia in TKI-sensitive *EGFR^L858R^* GEMM with concurrent *PIK3CA* mutant

Previous studies indicated that *PIK3CA* mutant drives resistance to TKIs in *EGFR*-mutant lung cancer^19,20^. We then assessed the effect of single-agent Osi on mice carrying TKI-sensitive *EGFR* mutation (*EGFR^L858R^*) plus *PIK3CA* mutation (Fig. 3A). Osi treatment exhibited potent tumor-inhibitory effects, resulting in nearly complete tumor regression supported by dramatic decrease of tumor burdens and tumor numbers (Fig. 3B-D). In parallel, we observed increased weights in the TA, QUAD, GAST skeletal muscles, as well as in WAT tissues, indicative of the notable alleviation of cachexia (Fig. 3E-F). Moreover, Osi treatment mitigated atrophic changes in these tissues, as evidenced by an increase in muscle fiber CSA and adipocyte size (Fig. 3G-K). These data suggest that, in the TKI-sensitive GEMM, tumor malignancy and cachexia mediated by the concurrent *PIK3CA* mutant could be effectively suppressed by Osi treatment.

**Fig. 3.**
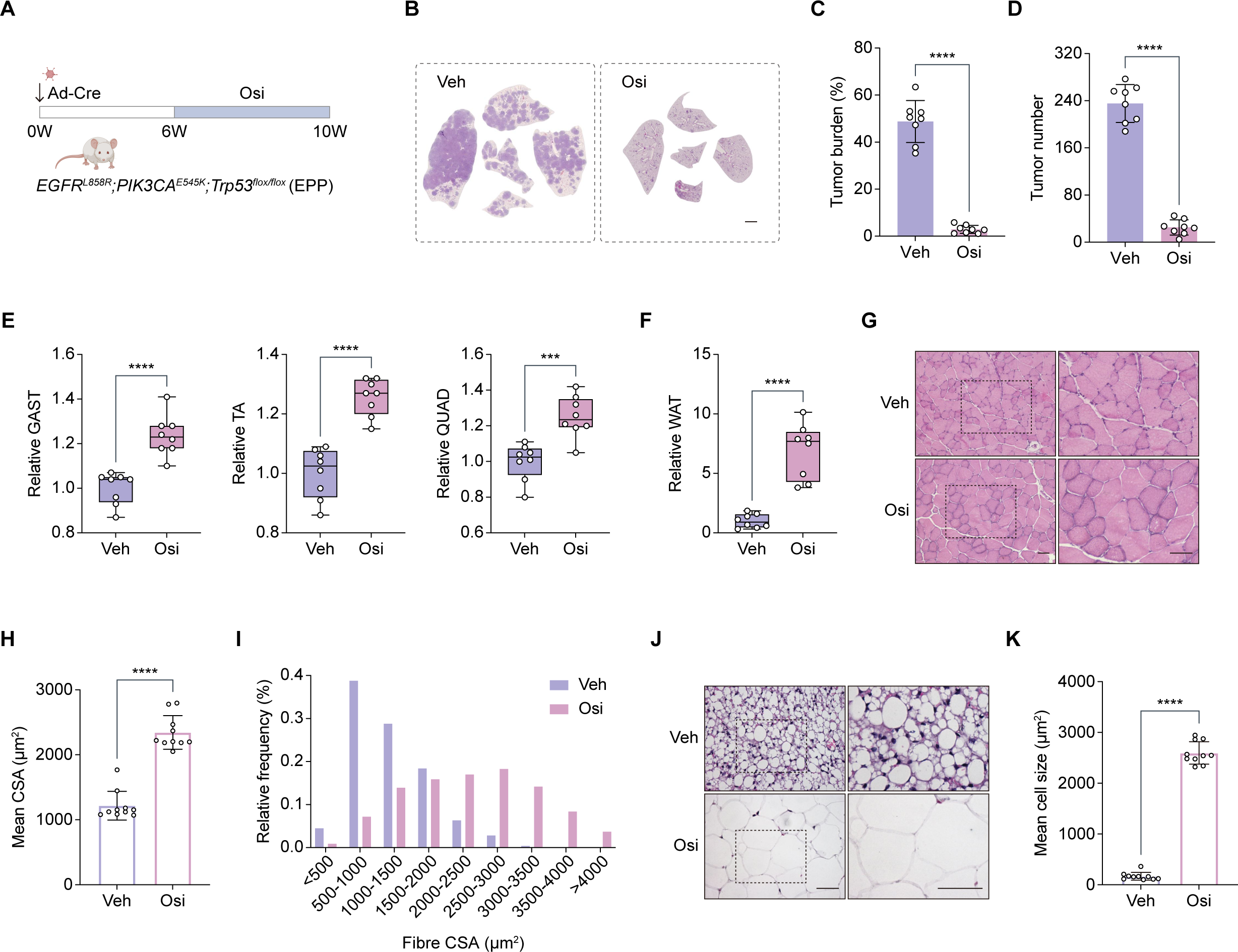
Osimertinib effectively suppresses cancer progression and alleviates cachexia in TKI-sensitive *EGFR*-mutant GEMM. A. Schematic diagram of osimertinib treatment strategy. B-D. Representative H&E staining images of lung tissue (B), statistical analyses of tumor burden (C) and tumor number (D) in EPP mice following gavage of either vehicle (Veh) (n=8) or 5mg/kg/day osimertinib (Osi) (n=8) for 28 days. Scale bar, 2 mm. E-F. Assessed weights of the gastrocnemius (GAST), tibialis anterior (TA), quadriceps (QUAD) (E), and white adipose tissue (WAT) (F) in EPP mice following gavage of either vehicle (Veh) (n=8) or 5mg/kg/day osimertinib (Osi) (n=8) for 28 days. Data were normalized based on the average values of corresponding tissue in mice receiving vehicle with the same gender. G-I. Representative micrographs (G), average fiber cross-sectional area (CSA) (H), and fiber CSA distribution (I) of gastrocnemius in EPP mice following gavage of either vehicle (Veh) or 5mg/kg/day osimertinib (Osi) for 28 days. Scale bar, 50 μm. J-K. Representative micrographs (J), and average size of adipocytes (K) of white adipose tissue in EPP mice following gavage of either vehicle (Veh) or 5mg/kg/day osimertinib (Osi) for 28 days. Scale bar, 50 μm. n=10 microscopic fields in each group for (H), (I) and (K). Data are presented as mean ± SD. ***P < 0.001, ****P< 0.0001 by two-tailed unpaired Student’s t test (C-F, H, K).

### Chemotherapy inhibits tumor growth but fails to alleviate cachexia in Osi-resistant *EGFR^TLCS^* GEMM with concurrent *PIK3CA* mutant

Once relapses from Osi therapy, chemotherapy, e.g., PEM and CDDP, is routinely used as the conventional therapeutic strategy for patients^18^. We next developed *EGFR^TLCS^*mice carrying *L858R*, *T790M*, and *C797S* mutations (Fig. S5A), which confer resistance to all three generations of TKIs used in clinical practice^17^. We generated the *EGFR^TLCS^;Trp53^flox/flox^*(TLCS-P) and *EGFR^TLCS^;PIK3CA^E545K^;Trp53^flox/flox^* (TLCS-PP) cohorts for further study, and found PI3K-AKT signaling was significantly enriched in mouse lung tumors derived from TLCS-PP mice (Fig. S5B). Compared to the other two groups, TLCS-PP mice showed an earlier onset of cachexia development, characterized by continuous body weight loss and reductions in both lean and fat mass (Fig. 4A-D).

**Fig 4.**
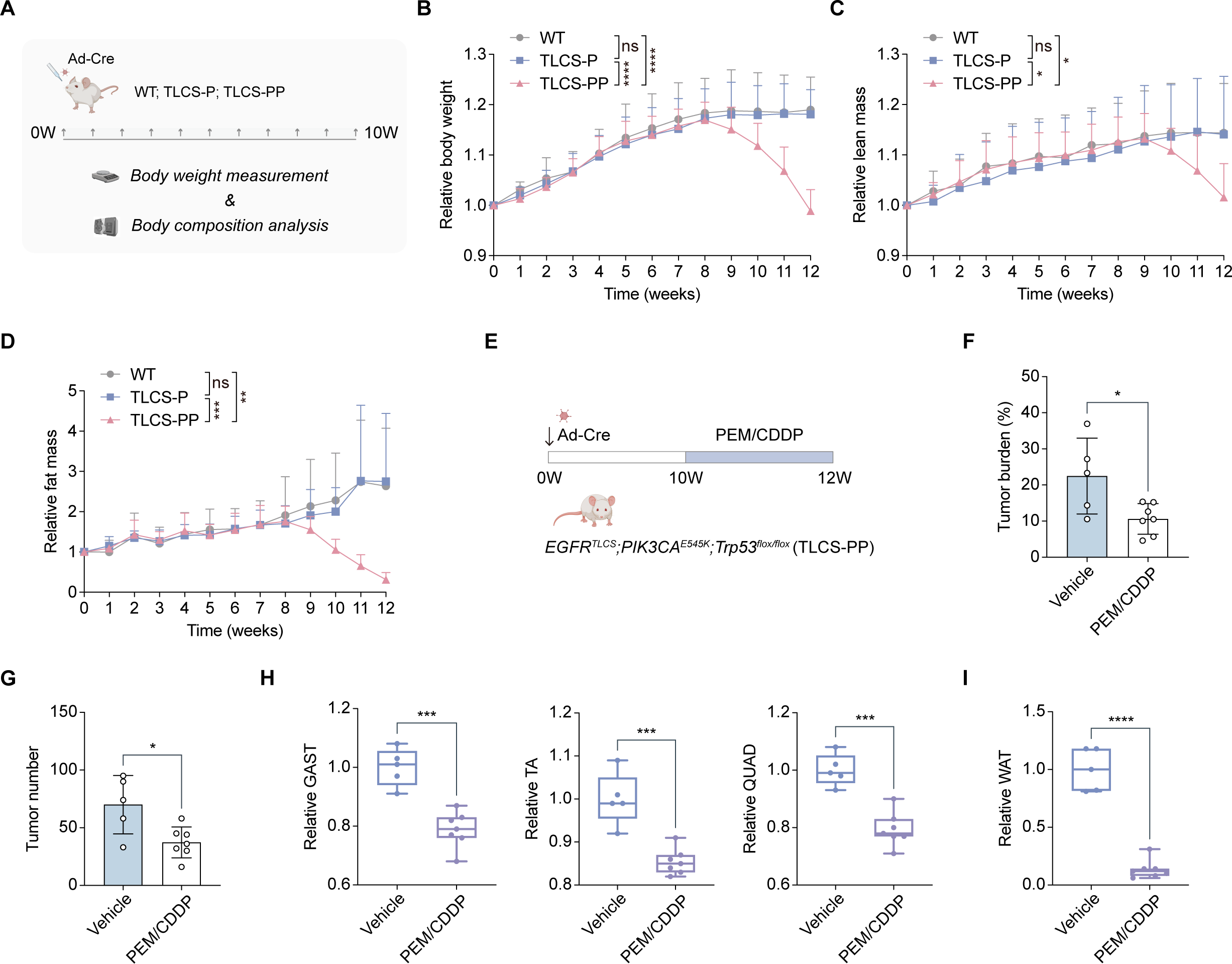
Chemotherapy fails to alleviate cachexia despite notable tumor suppression in TKI-resistant *EGFR*-mutant GEMM. A. Schematic diagram of long-term monitoring strategy of mouse body weight and body composition. B-D. Relative changes in body weight (B), lean mass (C), and fat mass (D) of WT mice (n=10), TLCS-P mice (n=10), and TLCS-PP mice (n=10) over time following intranasal delivery of Ad-Cre for 0-12 weeks. Data were normalized based on the values at the time of intranasal induction, and data collected at the 12-week post-induction were utilized for differential analysis. E. Schematic diagram of PEM/CDDP treatment strategy. F-G. Statistical analyses of tumor burden (F) and tumor number (G) in TLCS-PP mice following intraperitoneal injection of either vehicle (Veh) (n=5) or PEM/CDDP (n=7) for 14 days. H-I. Assessed weights of the gastrocnemius (GAST), tibialis anterior (TA), quadriceps (QUAD) (H), and white adipose tissue (WAT) (I) in TLCS-PP mice following intraperitoneal injection of either vehicle (Veh) (n=5) or PEM/CDDP (n=7) for 14 days. Data were normalized based on the average values of corresponding tissue in mice receiving vehicle with the same gender. Data are presented as mean ± SD. *P < 0.05, **P < 0.01, ***P < 0.001, ****P< 0.0001 by one-way ANOVA (B-D), two-tailed unpaired Student’s t test (F-I). ns: not significant.

We next treated the TLCS-PP mice with the standard second-line chemotherapy regimen of PEM and CDDP (Fig. 4E). We found chemotherapy resulted in a significant reduction of both tumor burdens and numbers, indicative of its effectiveness in suppressing tumor progression (Fig. 4F-G). Consistent with the adverse effects of chemotherapy on cachexia^42^, the PEM/CDDP combination failed to exert protective effect against cachexia development, as evidenced by the reduced weights of skeletal muscle and adipose tissue (Fig. 4H-I). These data suggest that the *PIK3CA* mutant-associated cachexia in Osi-resistant cases can’t be alleviated by chemotherapy despite notable tumor regression, emphasizing the need to explore the mechanisms underlying *PIK3CA* mutant in driving cachexia.

### *PIK3CA* mutant drives cachectic inflammation through NF-**κ**B activation

We then performed RNA-seq on mouse lung tumors derived from both Osi-sensitive and Osi-resistant GEMMs with or without *PIK3CA* mutant. Enrichment of epithelial-mesenchymal transition and inflammatory response pathways has been documented as a clinical feature of primary lung tumors from cachexic patients^30^. Interestingly, we observed the enrichment of both pathways in the EPP and TLCS-PP tumors compared to their respective controls (Fig. 5A, S5D-E). Moreover, the EPP and TLCS-PP tumors exhibited the enrichment of NF-κB signaling, which is known as the upstream regulator of multiple cachexia-associated pro-inflammatory factors^43^ (Fig. 5A, S5E). Upregulation of key factors in these tumors was further confirmed by RNA-seq and real-time PCR analyses (Fig. 5B-D, S5F).

**Fig. 5.**
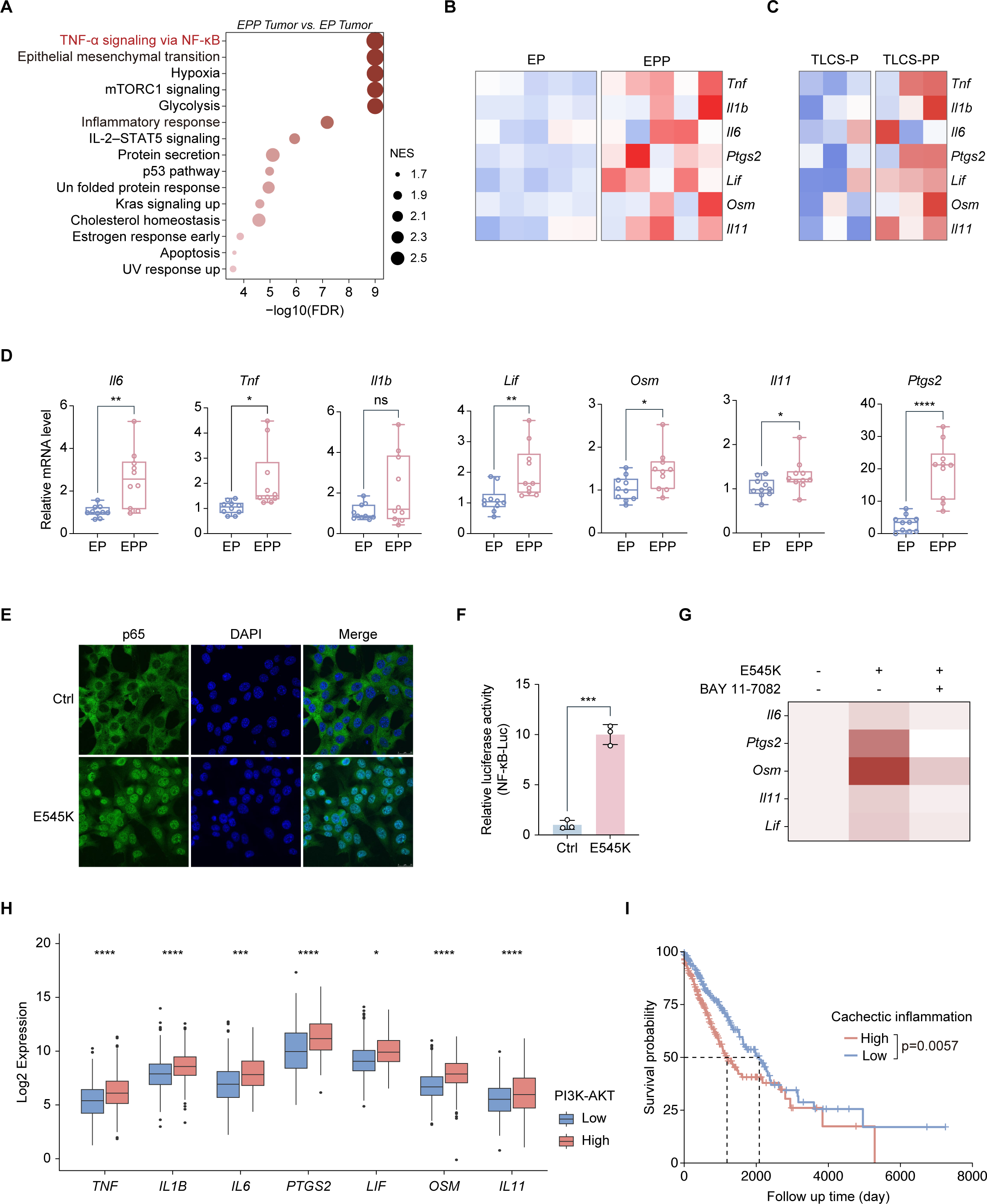
P*I*K3CA mutant increases cachexia-associated pro-inflammatory factors expression through NF-**κ**B activation. A. Dot plot shows Hallmark gene sets enriched in EPP tumors (n=6) versus EP tumors (n=6) based on GSEA analysis. The dot size represents the normalized enrichment score (NES). The dot color indicates enrichment significance. B-C. Gene expression heatmap of cachexia-associated pro-inflammatory factors in EP (n=6) and EPP (n=6) tumor tissues (B); as well as TLCS-P (n=3) and TLCS-PP (n=3) tumor tissues (C). D. Real-time PCR detection of mRNA levels for indicated genes in EP (n=10) and EPP (n=10) tumor tissues. E. Representative micrographs of immunofluorescence staining on EP cell line stably expressing empty vector (Ctrl) or *PIK3CA E545K* mutant (E545K). p65 in green, nucleus in blue (DAPI staining). Scale bar, 25 μm. F. HEK293T cells were co-transfected with NF-κB-Luc, Renilla plasmid, and either empty vector (Ctrl) or *PIK3CA E545K* mutant (E545K). Firefly luciferase activity was measured 48h after transfection. Firefly fluorescence were normalized based on the Renilla fluorescence values. G. Heat map displays the expression of indicated genes in the EP cell line stably expressing empty vector (Ctrl) or *PIK3CA E545K* mutant (E545K) with or without 10μM BAY 11L7082 treatment for 6 hours. H. Expression levels of indicated genes in non-small cell lung cancer samples with either high or low levels of PI3K-AKT signaling in The Cancer Genome Atlas (TCGA) database. I. Kaplan-Meier overall survival (OS) of non-small cell lung cancer patients with high or low cachectic inflammation signature in The Cancer Genome Atlas (TCGA) database. Data are presented as mean ± SD. *P < 0.05, **P < 0.01, ***P < 0.001, ****P< 0.0001 by two-tailed unpaired Student’s t test (D, F), multiple t test (H), Mantel–Cox test (I). ns: not significant.

To further clarify the role of NF-κB signaling in this process, we performed immunostaining analyses on the EP cells with ectopic expression of *PIK3CA* mutant (Fig. S2D). We found that *PIK3CA* mutant notably increased the nuclear translocation of p65 (Fig. 5E). The enhancement of NF-κB activity was further confirmed by luciferase detection (Fig. 5F). Additionally, cachexia-associated factors were found upregulated following ectopic expression of *PIK3CA* mutant and attenuated by the treatment with NF-κB inhibitor BAY 11-7082 (Fig. 5G).

We further analyzed NSCLC patient data from The Cancer Genome Atlas (TCGA) database. Increased expression levels of cachexia-associated factors were observed in samples exhibiting elevated PI3K-AKT signaling (Fig. 5H). We next consolidated these factors into a cachectic inflammation signature, and found that a higher cachectic inflammation signature was associated with shorter patient overall survival (Fig. 5I). These data indicate that *PIK3CA* mutant triggers NF-κB signaling, leading to the upregulation of cachexia-associated pro-inflammatory factors, which are associated with poor prognosis of lung cancer patients.

### Combined aspirin treatment attenuates cachexia in *EGFR* mutant lung cancer with concurrent *PIK3CA* mutant

Aspirin, a well-established non-steroidal anti-inflammatory drug (NSAID), has been reported to inhibit the activity of NF-κB signaling^44^. To assess the effect of aspirin on inflammation inhibition, we treated EP tumor cells with aspirin *in vitro* and found that aspirin effectively suppressed the pro-inflammatory factors upregulated by ectopic expression of *PIK3CA* mutant (Fig. S6A). We then treated the TLCS-PP mice with aspirin alone or combined with PEM/CDDP (Fig. 6A). Interestingly, aspirin treatment alone significantly mitigated weight loss without notable tumor regression (Fig. 6B-D, S6B). When combined aspirin with chemotherapy, we observed pronounced tumor regression, and the weight loss became similar to the vehicle group. Notably, we found that aspirin treatment led to increased weights in both skeletal muscle and adipose tissue, indicating effective suppression of cachexia (Fig. 6E-F). Aspirin also resulted in increased muscle fiber CSA and adipocyte size (Fig. S6C-F), histologically confirming the recovery from atrophic changes. These data suggest that aspirin, despite no tumor inhibition role, effectively mitigates cachexia progression of TKI-resistant lung cancer with concurrent *PIK3CA* mutant.

**Fig. 6.**
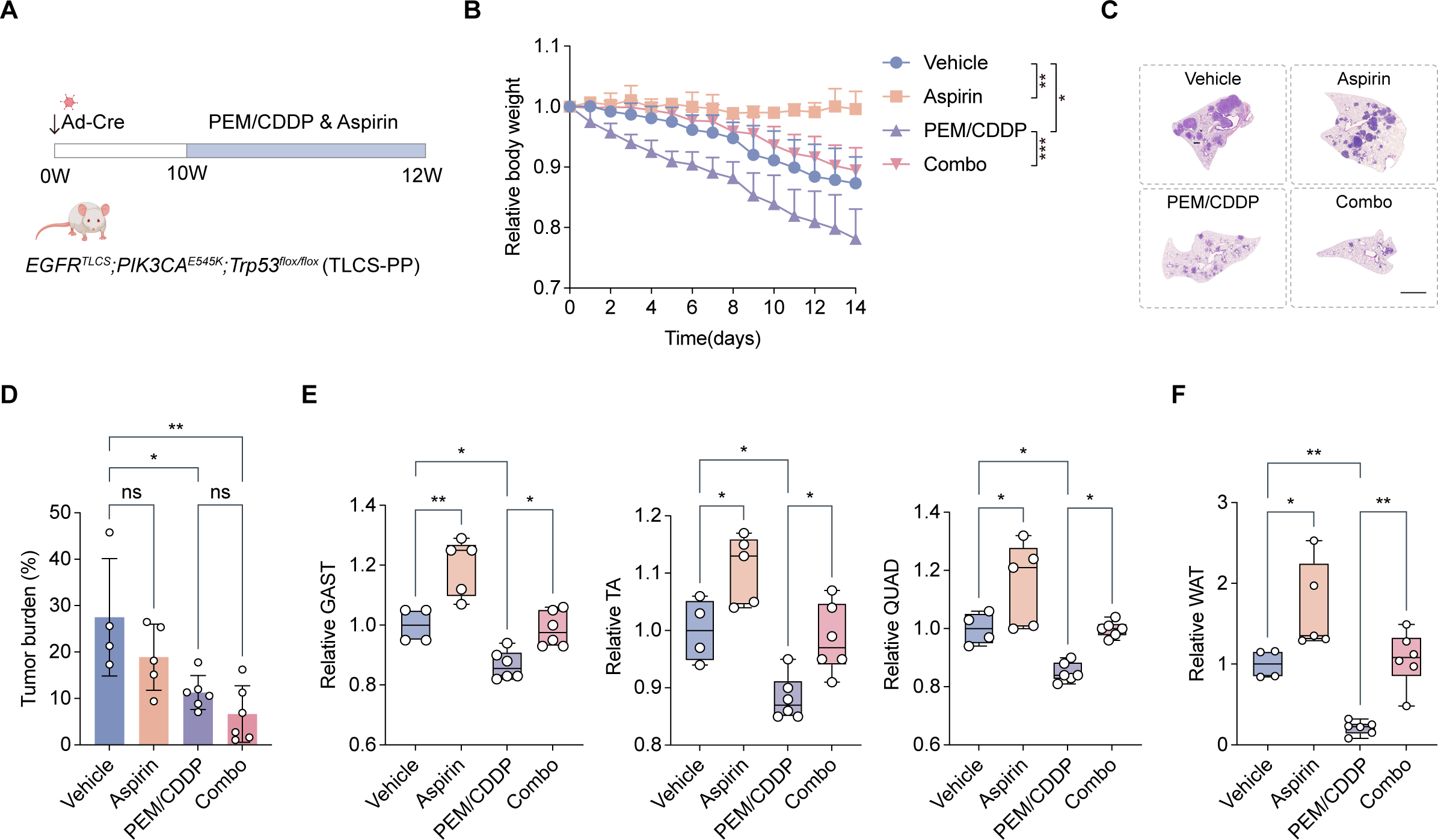
Aspirin ameliorates cachexia driven by *PIK3CA* mutant. A. Schematic diagram of combined treatment strategy involving PEM/CDDP and aspirin. B. Relative changes over time in body weight during intraperitoneal administration of either vehicle (Veh) (n=4), 50mg/kg aspirin (n=5), PEM/CDDP (n=6), or combined treatment (Combo) (n=6). Data were normalized based on the values at the onset of drug administration, and data collected at the 14-day time point following drug treatment were utilized for differential analysis. C-D. Representative H&E staining images of lung tissue (C) and statistical analyses of tumor burden (D) in TLCS-PP mice following intraperitoneal administration of either vehicle (Veh) (n=4), 50mg/kg aspirin (n=5), PEM/CDDP (n=6), or combined treatment (Combo) (n=6) for 14 days. Scale bar, 2 mm. E-F. Assessed weights of the gastrocnemius (GAST), tibialis anterior (TA), quadriceps (QUAD) (E), and white adipose tissue (WAT) (F) in TLCS-PP mice following intraperitoneal administration of either vehicle (Veh) (n=4), 50mg/kg aspirin (n=5), platinum-based doublet chemotherapy (PEM/CDDP) (n=6), or combined treatment (Combo) (n=6) for 14 days. Data were normalized based on the average values of corresponding tissue in mice receiving vehicle with the same gender. Data are presented as mean ± SD. *P < 0.05, **P < 0.01, ***P < 0.001 by one-way ANOVA (B, D-F). ns: not significant.

## Discussion

*PIK3CA* mutation has long been considered an important oncogenic driver in lung tumorigenesis, although it frequently co-occurs with *EGFR* mutation, *KRAS* mutation and *ALK* fusion^6,7^. Previous studies have convincingly shown that *EGFR* mutation, *KRAS* mutation or *ALK* fusion alone is sufficient to drive lung cancer initiation in mice^12-14^, and their co-expression even results in tumor growth inhibition^9,11^. This raises an interesting question about the real function of *PIK3CA* mutant in lung tumorigenesis. A previous study demonstrates that doxycycline-induced *PIK3CA* mutant transgene expression is sufficient to drive lung cancer initiation in mice^5^. In contrast, we find that Ad-Cre-mediated expression of *PIK3CA* mutant transgene plays a limited role in mouse lung tumor initiation. Consistently, Trejo CL *et al*. find no lung tumor formation in mice with endogenous expression of *PIK3CA* mutant^45^. No tumor is detectable in this model even after one year of *PIK3CA* mutant induction^46^. We reason that this discrepancy might be explained by various mouse models, the strength of PI3K pathway activation, and/or genetic backgrounds^45^. Nonetheless, clinical studies of multiple cancer types tend to support the incapability of *PIK3CA* mutant in driving tumor initiation. For example, human endometrial epithelium, which is pathologically normal, is found to harbor frequent *PIK3CA* mutations^47^. Similarly, *PIK3CA* mutations are detectable in pathologically normal human esophagus^48^. This study also reports no esophageal tumor detectable in *PIK3CA*-mutant transgenic mice^48^. These studies collectively demonstrate that *PIK3CA* mutant does not seem to contribute to cancer initiation as *EGFR* mutant, *KRAS* mutant, or *ALK* fusion does. Instead, it is more likely to play a secondary role in tumorigenesis, especially when concurrent with oncogenic drivers such as *EGFR* mutant, *KRAS* mutant, or *ALK* fusion. In line with this, we find that *PIK3CA* mutant mainly contributes to lung cancer malignant progression when coexisting with *EGFR* mutant, e.g., the concurrent *PIK3CA* mutant drives cachexia via activation of inflammatory signaling and leads to poor survival of mice bearing *EGFR*-mutant lung cancer. These findings provide a reasonable explanation for the longstanding paradox about the real biological function of concurrent *PIK3CA* mutant in lung tumorigenesis.

Another paradox exists in the seemingly contradictory impact of the *PIK3CA* mutant upon PFS and OS of *EGFR*-mutant lung cancer patients following TKI therapy^21,22^. Eng J *et al*. find that concurrent *PIK3CA* mutation is associated with shorter survival in *EGFR*-mutant lung cancer; nevertheless, they find no difference of the best objective response, time to best response, time to progression, or TKI duration time between patients with and without *PIK3CA* mutation^22^. Similarly, Song Z *et al*. find that *PIK3CA* mutations lead to shorter OS, while these mutations do not seem to affect the PFS of *EGFR*-mutant patients^21^. Previous studies have observed the acquisition of *PIK3CA* mutations in relapsed patients after TKI resistance, indicative of its potential contribution to drug resistance, despite occurring at a low rate^19,20^. However, these mutations do not seem to impact the efficacy of TKIs. For example, Wu SG *et al*. find that the rate of *PIK3CA* mutation is comparable in relapsed patients vs. treatment-naïve patients, with no significant impact on therapeutic response and PFS between *PIK3CA* mutation-positive and -negative cases^49^. Another study also reports that 22 out of 27 *EGFR*-mutant patients show partial responses, including those with *PIK3CA* co-mutations^50^. These clinical observations point to a potentially dispensable role of *PIK3CA* mutant as therapeutic target. Indeed, our mouse model findings support that *PIK3CA* mutant contributes to lung tumorigenesis by driving cachexia rather than cancer initiation. Since *PIK3CA* mutant is not important for tumor initiation, tumors are less likely to rely on these mutations for survival. This explains why Osi treatment effectively inhibits the growth of *EGFR*-mutant lung cancer even with concurrent *PIK3CA* mutations. However, once patients relapse from TKI therapy and the tumors develop TKI resistance, the impact of *PIK3CA* mutant begins to unleash, with cachexia driven by *PIK3CA* mutant gradually worsening and ultimately leading to poor patient prognosis. Our data from both TKI-sensitive and -resistant *EGFR*-mutant mouse models collectively support this view. We find that Osi effectively inhibits both tumor growth and cachexia progression in mice bearing tumors with *EGFR^L858R^* (TKI-sensitive) and *PIK3CA* co-mutations. Moreover, PIK3CA-associated cachexia is unresponsive to second-line chemotherapy, as demonstrated by the reduced tissue weights in mice bearing tumors with *EGFR^TLCS^* (TKI-resistance) and *PIK3CA* co-mutations. This is in line with clinical observations documenting the adverse effects of chemotherapy on cachexia progression^42^.

Systemic inflammation is recognized as the driving force of cachexia development in cancer^51^. We find that *PIK3CA* mutant activates NF-κB signaling, leading to increased expression of those cachexia-associated pro-inflammatory factors, such as IL-6 and LIF^25^. In addition to pro-inflammatory factors secreted by tumor and immune cells, chemotherapy is recognized as a significant source of systemic inflammation in the patients’ macroenvironment^52^. Recent reports demonstrate that chemotherapy induces the upregulation of pro-inflammatory cytokines and chemokines, contributing to the progression of cachexia^53^, further underscoring the significance of inflammation in cachexia progression.

Lastly, we find that aspirin, a type of NSAID, effectively inhibits the progression of cachexia. Although inflammation is widely recognized as a hallmark of cachexia development, previous clinical studies indicate that targeting TNF-α, IL-6, or IL-1 individually demonstrates limited effectiveness^54^. We reason that attributing cachexia progression to a single factor is challenging, as various pro-inflammatory factors interact through complex feedback loops^51^. Therefore, broader-spectrum anti-inflammatory drugs might offer a more effective intervention. Interestingly, previous studies show that colorectal cancer patients with *PIK3CA* mutations receiving regular aspirin treatment exhibit improved prognosis^55,56^. A multicenter, multinational, prospective randomized trial recently provides the first evidence of a positive effect of adjuvant aspirin in colon cancer patients with *PIK3CA* mutations^57^, further emphasizing the significance of adjuvant anti-inflammatory treatment in enhancing the prognosis of these patients. Our study provides new insights into the benefit of aspirin for *PIK3CA*-mutant patients with colon cancer. To validate our findings across other cancer types, we further perform Gene Set Enrichment Analysis (GSEA) on a recently published pre-clinical melanoma model^58^, and observed enrichment of PI3K-AKT signaling in xenografts capable of inducing cachexia (data not shown). Consistently, a previous study also reveals an association between *PIK3CA* mutant and weight loss in pancreatic ductal adenocarcinoma (PDAC) patients^59^. Given the significant contribution of *PIK3CA* mutant to poor prognosis by driving cachexia, it will be important to further investigate its broader implications in lung cancer and beyond.

## Materials and methods

### Mouse models

The *Trp53^flox/flox^* mice were originally provided by Dr. Tyler Jacks (Cambridge, MA). The transgenic mouse models, including *PIK3CA^E545K^*, *EGFR^L858R^*, and *EGFR^TLCS^*mice were generated using CRISPR/Cas9 technology. The brief process is as follows: homologous recombination vector (donor vector) is composed of a 5′ homology arm, the indicated coding sequence (CDS), and a 3′ homology arm. Cas9 mRNA, gRNA, and the donor vector were microinjected into the fertilized eggs of C57BL/6J mice to generate the F0 generation mice. All mice were kept in specific pathogen-free environment at Shanghai Institute of Biochemistry and Cell Biology and treated in strict accordance with protocols (SIBCB-2101008) approved by the Institutional Animal Care and Use Committee of the Shanghai Institutes for Biological Sciences, Chinese Academy of Sciences. Mice were delivered with Ad-Cre virus (2 × 10^6^ p.f.u.) via nasal inhalation at 6-8 weeks of age.

Osimertinib (MCE, HY-15772) was prepared in a solution containing 5% DMSO, 40% PEG300, 5% Tween-80 and 50% saline. Pemetrexed (MCE, HY-10820) or aspirin (MCE, HY-14654) was prepared in a solution containing 10% DMSO, 40% PEG300 and 50% saline. Cisplatin (MCE, HY-17394) was formulated in saline. For osimertinib treatment, *EGFR^L858R^;PIK3CA^E545K^;Trp53^flox/flox^* mice were gavage daily with osimertinib (5 mg/kg/day) 6 weeks after Ad-Cre infection. Control mice were administered the vehicle solution (5% DMSO: 40% PEG300: 5% Tween-80: 50% saline). For chemotherapy and combination therapy, *EGFR^TLCS^;PIK3CA^E545K^;Trp53^flox/flox^*mice were administered pemetrexed (50 mg/kg/day) combined with cisplatin (4 mg/kg/day) or aspirin (50 mg/kg/day) or both via intraperitoneal injection 10 weeks post Ad-Cre infection. Pemetrexed and cisplatin were administered weekly, and aspirin was given daily. Control mice were administered the vehicle solution (10% DMSO: 40% PEG300: 50% saline). All mice were sacrificed for gross inspection and histopathological examination. Tumor number, tumor burden, and tumor size were analyzed using ImageJ software.

### Cell culture and lentivirus infection

HEK-293T and EP cells were cultured in DMEM (HyClone) supplemented with 10% fetal bovine serum (FBS) and 1% penicillin/streptomycin (P/S). Cells were used for experiments within 10 to 20 passages from thawing. All cell lines were routinely tested for mycoplasma. For the establishment of stable overexpression cell line, HEK-293T cells were transfected with a 4:3:2 ratio of pCDH-CMV-PIK3CA E545K-EF1-Puro plasmid, psPAX2 plasmid (Addgene #12260), and pMD2.G plasmid (Addgene #12259). Lentiviral particles generated were then transduced into EP cells, followed by puromycin selection (3 μg/mL; Sigma-Aldrich) initiated 48 hours post-transfection and continued for an additional 2 days.

### Immunoblotting

Whole-cell lysates of cell lines or murine tumors were generated using lysis buffer (10% SDS, 1 mM DTT, and glycerin) and incubated at 100 °C for 10 min. Equal amounts of total protein were separated by SDS-PAGE and transferred onto PVDF membranes. Protein samples were probed with specific antibodies against p110α (CST, 4249, 1:1000), AKT (CST, 9272, 1:1000), p-AKT (CST, 4070, 1:1000), S6 (CST, 2217, 1:1000), p-S6 (CST, 2215, 1:1000) and β-actin (Abclonal, AC026, 1:50000). Protein expression was assessed by Pierce ECL Western Blotting Substrate (Thermo Fisher Scientific) and detected on SAGECREATION (Sage Creation Science Co.).

### RNA Isolation, reverse transcription, and real-time PCR

Total RNA from cultured cells or tissue samples was extracted using TRIzol (Invitrogen), and complementary DNA was synthesized from 1 μg of RNA using the PrimeScript RT Reagent Kit (TaKaRa). Real-time PCR was performed on a LightCycler^®^ 96 System (Roche) using SYBR Green I Master (Roche). β-actin was utilized as the internal control. The following

primers were used: *Il1b* F: 5’-GCAACTGTTCCTGAACTCAAC-3’, R: 5’-ATCTTTTGGGGTCCGTCAACT-3’. *Il6* F: 5’-TCTGCAAGAGACTTCCATCCAGTTGC-3’, R: 5’-AGCCTCCGACTTGTGAAGTGGT-3’. *Il11* F: 5’-CTGACGGAGATCACAGTCTGGA-3’, R: 5’-GGACATCAAGTCTACTCGAAGCC-3’. *Lif* F: 5’-AAAAGCTATGTGCGCCTAACA-3’, R: 5’-GTATGCGACCATCCGATACAG-3’. *Osm* F: 5’-ACGGTCCACTACAACACCAG-3’, R: 5’-CCATCGTCCCATTCCCTGAAG-3’. *Ptgs2* F: 5’-TTCAACACACTCTATCACTGGC-3’, R: 5’-AGAAGCGTTTGCGGTACTCAT-3’. *Tnf* F: 5’-CTGAACTTCGGGGTGATCGG-3’, R: 5’-GGCTTGTCACTCGAATTTTGAGA-3’. *Actb* F: 5’-GGCTGTATTCCCCTCCATCG-3’, R: 5’-CCAGTTGGTAACAATGCCATGT-3’.

### Soft agar colony formation assay

Soft agar assay was performed as previously described^60^. In detail, a bottom layer of 1% agar with complete medium was first solidified. An upper layer was then added, containing 500 cells suspended in a 0.4% agar medium mixture in 6-well plates. After 2–3 weeks of incubation, cells were stained with 0.005% crystal violet, and the number of colonies was counted.

### Immunofluorescence analyses

Cells grown on glass coverslips were washed with cold PBS and fixed with 4% paraformaldehyde (PFA) in PBS for 15 min at room temperature. Fixed cells were permeabilized with 0.2% Triton X-100 in PBS for 15 min. After blocking with 4% bovine serum albumin (BSA) in PBS for 1 h, cells were probed with p65 (CST, 8242, 1:500) overnight at 4°C. After washing three times with PBST, secondary antibodies were added and incubated for 1 h at room temperature. After washed with PBST, the coverslips were mounted onto glass slides using fluorescent mounting medium. Confocal images were captured using a Leica TCS SP8 system with a HC PL APO CS2 63×/1.40 oil objective.

### Luciferase reporter gene assay

HEK293T cells were seeded in 12-well plate at 3x10^5^ cells/well. 24 h after seeding, cells were transiently transfected with 1 μg of either *PIK3CA E545K* plasmid or the control vector, along with 1 μg NF-κB-Luc plasmid and 500 ng Renilla plasmid. 48 hours post-transfection, cells were harvested, and the Dual-Luciferase Reporter Assay System (Promega) was used for detection.

### Tissue collection

Mice were euthanized, and their body weight was recorded. The lungs with tumors were excised and fixed in 4% formalin; a portion of the tumors was snap-frozen for further analysis. Gastrocnemius (GAST), tibialis anterior (TA), quadriceps (QUAD), epididymal white adipose tissue (eWAT), and gonadal white adipose tissue (gWAT) were dissected and weighed. The white adipose tissues were fixed in 4% formalin for histological examination. One part of the skeletal muscle was snap-frozen for RNA-seq, while the remaining portion was embedded in optimal cutting temperature (OCT) compound for rapid freezing and subsequent histological examination.

### Body composition analysis and fat imaging

Mice were weighed and body composition was measured using Bruker’s minispec LF50 Body Composition Analyzer. The instrument provides measurements of three related components: fat, free body fluids, and lean tissue mass. The imaging of mouse adipose tissue was performed using the NM42-060H-I (Niumag) small animal magnetic resonance imaging (MRI) scanner.

### Metabolic cage

Mice were individually housed in CLAMS metabolic cages (Columbus Instruments) for a duration of 3 days. Oxygen consumption (VO2) and carbon dioxide expiration (VCO2) were measured for 1 min with 14 min intervals at a flow rate set at 0.72 liter per minute. Respiratory exchange ratio (RER) was calculated as the ratio of VCO2 to VO2. Simultaneously, locomotor activity and energy expenditure were recorded using the built-in detection system, while food intake was manually measured at the same designated time each day.

### RNA-seq analyses

Raw fastq data from RNA-seq were processed with Trimmomatic^61^ (v0.39) for adapter trimming and low-quality read filtering. The processed data were then aligned to the mm10 reference genome using STAR^62^ (v2.5.2b). Genes with zero expression in more than 70% of the samples were filtered out. FPKM normalization and log2 transformation were applied to the raw count data, followed by differential expression analysis using limma^63^ to identify differentially expressed genes (DEGs) between conditions. Pathway enrichment analysis was conducted using the Enrichr^64^ method based on the resulting DEGs. In addition, GSEA enrichment analysis was performed with the clusterProfiler^65^ package based on ranked genes.

### Statistical analyses

For comparing means of two groups, differences were analyzed by Student’s t test (two-tailed) and performed by Prism GraphPad software. For comparing means of three or more than three groups, differences among groups were analyzed by one-way ANOVA performed by Prism GraphPad software. P value <0.05 was considered statistically significant. Error bars were represented with SD. Survival analysis was performed using the Kaplan–Meier method.

## Acknowledgments

We thank Dr. Tyler Jacks and Dr. Liang Chen for providing the mice. This work was supported by the National Key Research and Development Program of China (grants 2022YFA1103900 to H.J.; 2020YFA0803300 to H.J.); the National Natural Science Foundation of China (grants 82303039 to Z.Q., 82341002 to H.J., 32293192 to H.J., 82030083 to H.J.); the Shanghai Sailing Program(23YF1452900 to Z.Q.); the Basic Frontier Scientific Research Program of Chinese Academy of Science (ZDBS-LY-SM006 to H.J.); the Innovative research team of high-level local universities in Shanghai (SSMU-ZLCX20180500 to H.J.).

## Author Contributions

H.J. conceived the idea and designed the experiments. M.Y. and Z.Q. performed all experiments and analyzed the data. S.T. and X.C. performed the bioinformatics analyses. Y.Z. established primary cell line. L.C. and L.C. provided technical assistance and helpful comments. H.J. and M.Y. wrote the manuscript. All authors approved the final version.

## Illustration Tool

The graphical abstract image is created with BioRender.

---

**Fig. S1 *PIK3CA* mutant shows limited tumor initiation capacity in lung cancer.**

A. Schematic diagram of LoxP-stop-LoxP-*PIK3CA^E545K^* mouse model.

B. Representative H&E staining images of lung tissue from *PIK3CA^E545K^* mice following 40 weeks post Ad-Cre intranasal delivery. Scale bar, 2 mm.

C. Representative micrographs of *PIK3CA^E545K^* mice following 40 weeks post Ad-Cre intranasal delivery. Scale bar, 50 μm.

D. Mutation status and co-occurrence analysis of *PIK3CA* and *TP53* in non-small cell lung cancer patients from the *cBioPortal* cancer genomics database.

E. Representative H&E staining images of lung tissue from *PIK3CA^E545K^;Trp53^flox/flox^* mice following 40 weeks post Ad-Cre intranasal delivery. Scale bar, 2 mm.

F. Representative micrographs of *PIK3CA^E545K^;Trp53^flox/flox^* mice following 40 weeks post Ad-Cre intranasal delivery. Scale bar, 50 μm.

**Fig. S2 *PIK3CA* mutant promotes murine lung cancer malignant progression.**

A. Mutation status and co-occurrence analysis of *PIK3CA, EGFR, KRAS, BRAF, and MAP2K1* in non-small cell lung cancer patients from the *cBioPortal* cancer genomics database.

B. Schematic diagram of LoxP-stop-LoxP-*EGFR^L858R^* mouse model.

C. Schematic diagram of EP cell line establishment.

D. Immunoblotting analysis of phosphorylated AKT and phosphorylated S6 in EP cell line.

E-G, Representative images of soft agar colony formation (E), statistical analyses of colony numbers (F), and average colony size (G) of EP cell line stably expressing empty vector (Ctrl) or *PIK3CA E545K* mutant (E545K). Scale bar, 500 μm.

H. Immunoblotting analysis of phosphorylated AKT and phosphorylated S6 in *EGFR^L858R^; Trp53^-/-^* (EP) and *EGFR^L858R^; PIK3CA^E545K^; Trp53^-/-^* (EPP) *in situ* tumor tissues.

I-J. Statistical analyses of *in situ* tumor burden (I) and tumor number (J) in EP mice and EPP mice following 6 weeks (n=3), 8 weeks (n=3), and 10 weeks (n=5) post Ad-Cre intranasal delivery.

K. Statistical analyses of individual size of EP tumor (n=160) and EPP tumor (n=178) following 10 weeks post Ad-Cre intranasal delivery.

L. Statistical analyses of *in situ* tumor grade in EP mice (n=5) and EPP mice (n=5) following 10 weeks post Ad-Cre intranasal delivery.

M. Representative micrographs of *in situ* tumors with different grading characteristics in EP and EPP mice following 10 weeks post Ad-Cre intranasal delivery. Scale bar, 50 μm.

N. Kaplan-Meier curve shows the overall survival of EP mice (n=6), and EPP mice (n=6) receiving Ad-Cre delivery.

Data are presented as mean ± SD. *P < 0.05, **P < 0.01, ***P < 0.001, ****P< 0.0001 by two-tailed unpaired Student’s t test (F, G, K), multiple t test (I-J). ns: not significant.

**Fig. S3 *PIK3CA* mutant promotes lung cancer cachexia progression.**

A-B. Relative changes in lean mass (A), and fat mass (B) of WT mice (n=12), EP mice (n=12), and EPP mice (n=12) over time following intranasal delivery of Ad-Cre for 0-10 weeks. Data were normalized based on the values at the time of intranasal induction, and data collected at the 10-week post-induction were utilized for differential analysis.

C-E. Representative micrographs (C), average fiber cross-sectional area (CSA) (D), and fiber CSA distribution (E) of gastrocnemius tissue in WT mice, EP mice, and EPP mice following 8 weeks post Ad-Cre intranasal delivery. Scale bar, 50 μm.

F-G. Representative micrographs (F), and average size of adipocytes (G) of white adipose tissue in WT mice, EP mice, and EPP mice following 8 weeks post Ad-Cre intranasal delivery. Scale bar, 50 μm.

H-J. Statistical analyses of activity counts (H), energy expenditure (EE) (I), and respiratory exchange ratio (RER) (J) of WT mice (n=8), EP mice (n=8), and EPP mice (n=6) following 8 weeks post Ad-Cre intranasal delivery.

n=10 microscopic fields in each group for (D), (E) and (G). Data are presented as mean ± SD.

*P < 0.05, **P < 0.01, ***P < 0.001, ****P< 0.0001 by one-way ANOVA (A-B, E, G, H-J).

**Fig. S4 PI3K-AKT activation is associated with cachexia development in lung cancer patients.**

A. GSEA enrichment plot of PI3K-AKT signaling pathway in low-muscularity (LM) patients compared to high-muscularity (HM) patients in Cury *et al*. cohort.

B. Dot plot shows enriched KEGG pathways in the serum proteomics of cachexia patients compared to non-cachexia patients in Wang *et al*. cohort.

**Fig. S5 Gene expression characterization of murine tumors with *PIK3CA* mutations.**

A. Left part shows the schematic diagram of LoxP-stop-LoxP-*EGFR^TLCS^* mouse model; right part shows the representative H&E staining images of lung tissue from *EGFR^TLCS^*mice post Ad-Cre intranasal delivery. Scale bar, 2 mm.

B-E. GSEA enrichment plot of PI3K-AKT signaling pathway (B), inflammatory response (C), epithelial mesenchymal transition (D), and Tnf-α signaling via NF-κB pathway (E) in TCLS-PP tumors compared to TLCS-P tumors.

F. Real-time PCR detection of mRNA levels for indicated genes in TLCS-P (n=12) and TLCS-PP (n=12) tumor tissues.

Data are presented as mean ± SD. *P < 0.05, **P < 0.01, ***P < 0.001 by two-tailed unpaired Student’s t test (F).

**Fig. S6 Aspirin effectively ameliorates cachexia in *PIK3CA*-mutant lung cancer.**

A. Heat map of indicated genes expression in EP cell line stably expressing empty vector (Ctrl) or *PIK3CA E545K* mutant (E545K) with or without 2mM aspirin treatment for 24 hours.

B. Statistical analyses of tumor number in TLCS-PP mice following intraperitoneal administration of either vehicle (Veh) (n=4), 50mg/kg aspirin (n=5), PEM/CDDP (n=6), or combined treatment (Combo) (n=6) for 14 days.

C-D. Representative micrographs (C), and average fiber cross-sectional area (CSA) (D) of gastrocnemius tissue in TLCS-PP mice following intraperitoneal administration of either vehicle (Veh), 50mg/kg aspirin, PEM/CDDP, or combined treatment (Combo) for 14 days. Scale bar, 50 μm.

E-F. Representative micrographs (E), and average size of adipocytes (F) of white adipose tissue in TLCS-PP mice following intraperitoneal administration of either vehicle (Veh), 50mg/kg aspirin, PEM/CDDP, or combined treatment (Combo) for 14 days. Scale bar, 50 μm.

n=10 microscopic fields in each group for (D) and (F). Data are presented as mean ± SD. *P < 0.05, ****P< 0.0001 by one-way ANOVA (B, D, F). ns: not significant.

## References

1. Cancer Genome Atlas Research, N. Comprehensive molecular profiling of lung adenocarcinoma. Nature 511, 543-50 (2014).

2. Cancer Genome Atlas Research, N. Comprehensive genomic characterization of squamous cell lung cancers. Nature 489, 519-25 (2012).

3. Arafeh, R. & Samuels, Y. PIK3CA in cancer: The past 30 years. Semin Cancer Biol 59, 36–49 (2019).

4. Hoxhaj, G. & Manning, B.D. The PI3K-AKT network at the interface of oncogenic signalling and cancer metabolism. Nat Rev Cancer 20, 74–88 (2020).

5. Engelman, J.A. et al. Effective use of PI3K and MEK inhibitors to treat mutant Kras G12D and PIK3CA H1047R murine lung cancers. Nat Med 14, 1351–6 (2008).

6. Chaft, J.E. et al. Coexistence of PIK3CA and other oncogene mutations in lung adenocarcinoma-rationale for comprehensive mutation profiling. Mol Cancer Ther 11, 485–91 (2012).

7. Wang, L. et al. PIK3CA mutations frequently coexist with EGFR/KRAS mutations in non-small cell lung cancer and suggest poor prognosis in EGFR/KRAS wildtype subgroup. PLoS One 9, e88291 (2014).

8. Skoulidis, F. & Heymach, J.V. Co-occurring genomic alterations in non-small-cell lung cancer biology and therapy. Nat Rev Cancer 19, 495–509 (2019).

9. Unni, A.M., Lockwood, W.W., Zejnullahu, K., Lee-Lin, S.Q. & Varmus, H. Evidence that synthetic lethality underlies the mutual exclusivity of oncogenic KRAS and EGFR mutations in lung adenocarcinoma. Elife 4, e06907 (2015).

10. Tang, S. et al. Counteracting lineage-specific transcription factor network finely tunes lung adeno-to-squamous transdifferentiation through remodeling tumor immune microenvironment. Natl Sci Rev 10, nwad028 (2023).

11. Ambrogio, C., Barbacid, M. & Santamaria, D. In vivo oncogenic conflict triggered by co-existing KRAS and EGFR activating mutations in lung adenocarcinoma. Oncogene 36, 2309–2318 (2017).

12. Foggetti, G. et al. Genetic Determinants of EGFR-Driven Lung Cancer Growth and Therapeutic Response In Vivo. Cancer Discov 11, 1736–1753 (2021).

13. Ji, H. et al. LKB1 modulates lung cancer differentiation and metastasis. Nature 448, 807–10 (2007).

14. Qin, Z. et al. EML4-ALK fusions drive lung adeno-to-squamous transition through JAK-STAT activation. J Exp Med 221(2024).

15. Fu, K., Xie, F., Wang, F. & Fu, L. Therapeutic strategies for EGFR-mutated non-small cell lung cancer patients with osimertinib resistance. J Hematol Oncol 15, 173 (2022).

16. Thress, K.S. et al. Acquired EGFR C797S mutation mediates resistance to AZD9291 in non-small cell lung cancer harboring EGFR T790M. Nat Med 21, 560–2 (2015).

17. Herbst, R.S., Morgensztern, D. & Boshoff, C. The biology and management of non-small cell lung cancer. Nature 553, 446–454 (2018).

18. Tan, A.C. & Tan, D.S.W. Targeted Therapies for Lung Cancer Patients With Oncogenic Driver Molecular Alterations. J Clin Oncol 40, 611–625 (2022).

19. Le, X. et al. Landscape of EGFR-Dependent and -Independent Resistance Mechanisms to Osimertinib and Continuation Therapy Beyond Progression in EGFR-Mutant NSCLC. Clin Cancer Res 24, 6195–6203 (2018).

20. Chmielecki, J. et al. Analysis of acquired resistance mechanisms to osimertinib in patients with EGFR-mutated advanced non-small cell lung cancer from the AURA3 trial. Nat Commun 14, 1071 (2023).

21. Song, Z., Yu, X. & Zhang, Y. Mutation and prognostic analyses of PIK3CA in patients with completely resected lung adenocarcinoma. Cancer Med 5, 2694–2700 (2016).

22. Eng, J. et al. Impact of Concurrent PIK3CA Mutations on Response to EGFR Tyrosine Kinase Inhibition in EGFR-Mutant Lung Cancers and on Prognosis in Oncogene-Driven Lung Adenocarcinomas. J Thorac Oncol 10, 1713–9 (2015).

23. Ferrer, M. et al. Cachexia: A systemic consequence of progressive, unresolved disease. Cell 186, 1824–1845 (2023).

24. Argiles, J.M., Lopez-Soriano, F.J., Stemmler, B. & Busquets, S. Cancer-associated cachexia -understanding the tumour macroenvironment and microenvironment to improve management. Nat Rev Clin Oncol 20, 250–264 (2023).

25. Baracos, V.E., Martin, L., Korc, M., Guttridge, D.C. & Fearon, K.C.H. Cancer-associated cachexia. Nat Rev Dis Primers 4, 17105 (2018).

26. Bye, A. et al. Muscle mass and association to quality of life in non-small cell lung cancer patients. J Cachexia Sarcopenia Muscle 8, 759–767 (2017).

27. Ross, P.J. et al. Do patients with weight loss have a worse outcome when undergoing chemotherapy for lung cancers? Br J Cancer 90, 1905–11 (2004).

28. de Jong, C. et al. The association between skeletal muscle measures and chemotherapy-induced toxicity in non-small cell lung cancer patients. J Cachexia Sarcopenia Muscle 13, 1554–1564 (2022).

29. Jin, J., Visina, J., Burns, T.F., Diergaarde, B. & Stabile, L.P. Male sex and pretreatment weight loss are associated with poor outcome in patients with advanced non-small cell lung cancer treated with immunotherapy: a retrospective study. Sci Rep 13, 17047 (2023).

30. Al-Sawaf, O. et al. Body composition and lung cancer-associated cachexia in TRACERx. Nat Med 29, 846–858 (2023).

31. Oswalt, C. et al. Associations between body mass index, weight loss and overall survival in patients with advanced lung cancer. J Cachexia Sarcopenia Muscle 13, 2650–2660 (2022).

32. Yang, M., Shen, Y., Tan, L. & Li, W. Prognostic Value of Sarcopenia in Lung Cancer: A Systematic Review and Meta-analysis. Chest 156, 101–111 (2019).

33. Mytelka, D.S., Li, L. & Benoit, K. Post-diagnosis weight loss as a prognostic factor in non-small cell lung cancer. J Cachexia Sarcopenia Muscle 9, 86–92 (2018).

34. Frese, K.K. & Tuveson, D.A. Maximizing mouse cancer models. Nat Rev Cancer 7, 645–58 (2007).

35. DuPage, M., Dooley, A.L. & Jacks, T. Conditional mouse lung cancer models using adenoviral or lentiviral delivery of Cre recombinase. Nat Protoc 4, 1064–72 (2009).

36. Fearon, K. et al. Definition and classification of cancer cachexia: an international consensus. Lancet Oncol 12, 489–95 (2011).

37. Cury, S.S. et al. Low muscle mass in lung cancer is associated with an inflammatory and immunosuppressive tumor microenvironment. J Transl Med 21, 116 (2023).

38. Wang, D. et al. LCN2 secreted by tissue-infiltrating neutrophils induces the ferroptosis and wasting of adipose and muscle tissues in lung cancer cachexia. J Hematol Oncol 16, 30 (2023).

39. Fearon, K.C., Glass, D.J. & Guttridge, D.C. Cancer cachexia: mediators, signaling, and metabolic pathways. Cell Metab 16, 153–66 (2012).

40. Judge, S.M. et al. Skeletal Muscle Fibrosis in Pancreatic Cancer Patients with Respect to Survival. JNCI Cancer Spectr 2, pky043 (2018).

41. Talbert, E.E. et al. Modeling Human Cancer-induced Cachexia. Cell Rep 28, 1612–1622 e4 (2019).

42. Kimura, M. et al. Prognostic impact of cancer cachexia in patients with advanced non-small cell lung cancer. Support Care Cancer 23, 1699–708 (2015).

43. Pahl, H.L. Activators and target genes of Rel/NF-κB transcription factors. Oncogene 18, 6853–6866 (1999).

44. Kopp, E. & Ghosh, S. Inhibition of NF-kappa B by sodium salicylate and aspirin. Science 265, 956–9 (1994).

45. Trejo, C.L. et al. Mutationally activated PIK3CA(H1047R) cooperates with BRAF(V600E) to promote lung cancer progression. Cancer Res 73, 6448–61 (2013).

46. Green, S., Trejo, C.L. & McMahon, M. PIK3CA(H1047R) Accelerates and Enhances KRAS(G12D)-Driven Lung Tumorigenesis. Cancer Res 75, 5378–91 (2015).

47. Moore, L. et al. The mutational landscape of normal human endometrial epithelium. Nature 580, 640–646 (2020).

48. Herms, A. et al. Organismal metabolism regulates the expansion of oncogenic PIK3CA mutant clones in normal esophagus. Nat Genet (2024).

49. Wu, S.G., Chang, Y.L., Yu, C.J., Yang, P.C. & Shih, J.Y. The Role of PIK3CA Mutations among Lung Adenocarcinoma Patients with Primary and Acquired Resistance to EGFR Tyrosine Kinase Inhibition. Sci Rep 6, 35249 (2016).

50. Endoh, H., Yatabe, Y., Kosaka, T., Kuwano, H. & Mitsudomi, T. PTEN and PIK3CA expression is associated with prolonged survival after gefitinib treatment in EGFR-mutated lung cancer patients. J Thorac Oncol 1, 629–34 (2006).

51. Baazim, H., Antonio-Herrera, L. & Bergthaler, A. The interplay of immunology and cachexia in infection and cancer. Nat Rev Immunol 22, 309–321 (2022).

52. Swanton, C. et al. Embracing cancer complexity: Hallmarks of systemic disease. Cell 187, 1589–1616 (2024).

53. Englund, D.A., et al. Senotherapeutic drug treatment ameliorates chemotherapy-induced cachexia. JCI Insight 9 (2024).

54. Yue, M., Qin, Z., Hu, L. & Ji, H. Understanding cachexia and its impact on lung cancer and beyond. Chin Med J Pulm Crit Care Med 2, 95–105 (2024).

55. Domingo, E. et al. Evaluation of PIK3CA mutation as a predictor of benefit from nonsteroidal anti-inflammatory drug therapy in colorectal cancer. J Clin Oncol 31, 4297–305 (2013).

56. Liao, X. et al. Aspirin use, tumor PIK3CA mutation, and colorectal-cancer survival. N Engl J Med 367, 1596–606 (2012).

57. Güller, U. et al. 512O Adjuvant aspirin treatment in PIK3CA mutated colon cancer patients: The phase III, prospective-randomized placebo-controlled multicenter SAKK 41/13 trial. Annals of Oncology 35, S432 (2024).

58. Graca, F.A. et al. Progressive development of melanoma-induced cachexia differentially impacts organ systems in mice. Cell Rep 42, 111934 (2023).

59. Narasimhan, A. et al. Identification of Potential Serum Protein Biomarkers and Pathways for Pancreatic Cancer Cachexia Using an Aptamer-Based Discovery Platform. Cancers (Basel) 12(2020).

60. Qin, Z. et al. Phase separation of EML4-ALK in firing downstream signaling and promoting lung tumorigenesis. Cell Discov 7, 33 (2021).

61. Bolger, A.M., Lohse, M. & Usadel, B. Trimmomatic: a flexible trimmer for Illumina sequence data. Bioinformatics 30, 2114–20 (2014).

62. Dobin, A. et al. STAR: ultrafast universal RNA-seq aligner. Bioinformatics 29, 15–21 (2013).

63. Ritchie, M.E. et al. limma powers differential expression analyses for RNA-sequencing and microarray studies. Nucleic Acids Res 43, e47 (2015).

64. Chen, E.Y. et al. Enrichr: interactive and collaborative HTML5 gene list enrichment analysis tool. BMC Bioinformatics 14, 128 (2013).

65. Wu, T. et al. clusterProfiler 4.0: A universal enrichment tool for interpreting omics data. Innovation (Camb) 2, 100141 (2021).

