## Supplemental Figure 1-6 for "Concurrent *PIK3CA* mutant drives cachexia through inflammatory signaling in *EGFR* mutant lung cancer"

#### Figure S1

A

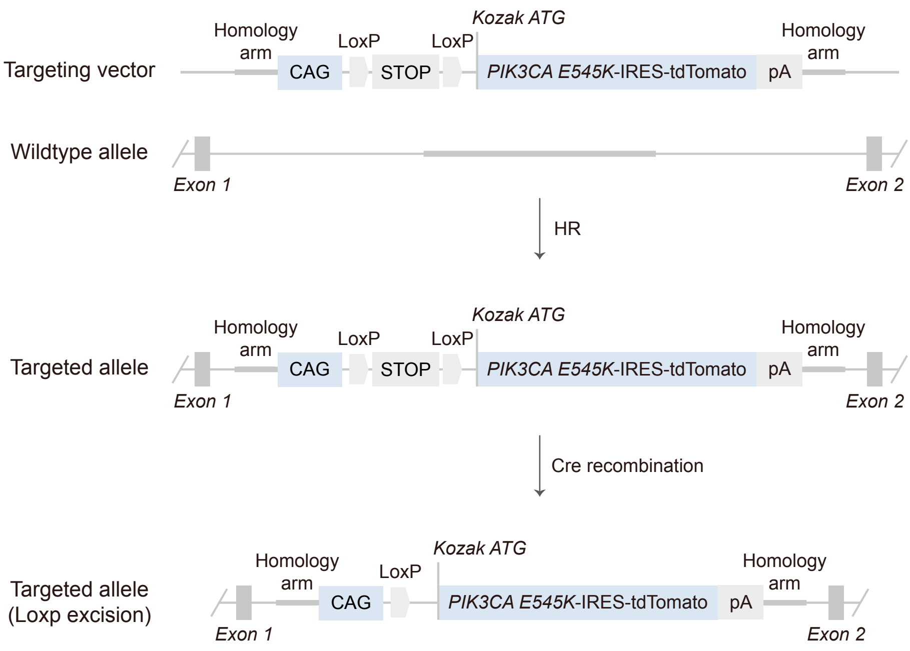


B

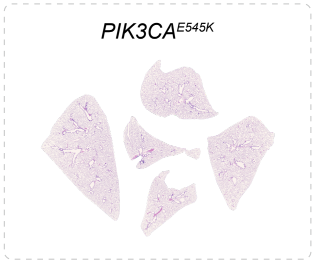


C

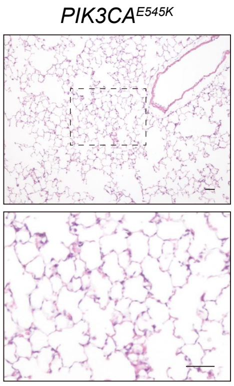


D

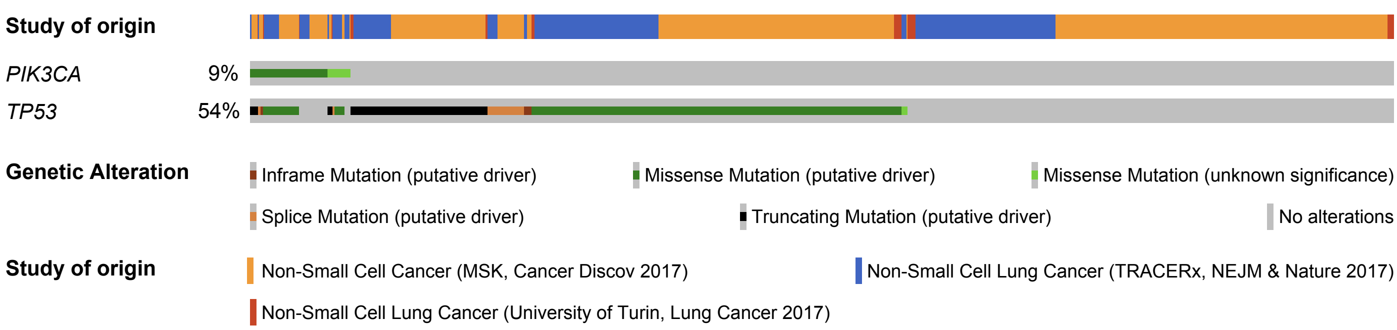


| a | b | Neither | a Not b | b Not a | Both | Log2 Odds Ratio | p-Value | q-Value | Tendency |
|---|---|---|---|---|---|---|---|---|---|
| *PIK3CA* | *TP53* | 597 | 42 | 683 | 81 | 0.753 | 0.008 | 0.008 | **Co-occurrence** |

E

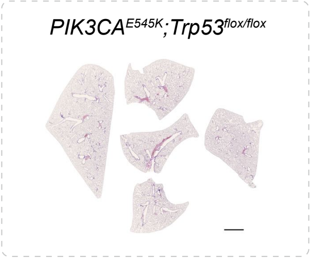


F

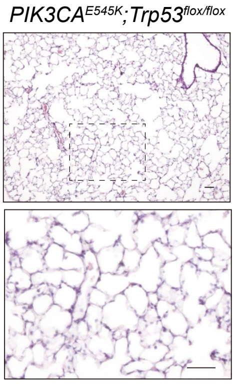

### Figure S2

A

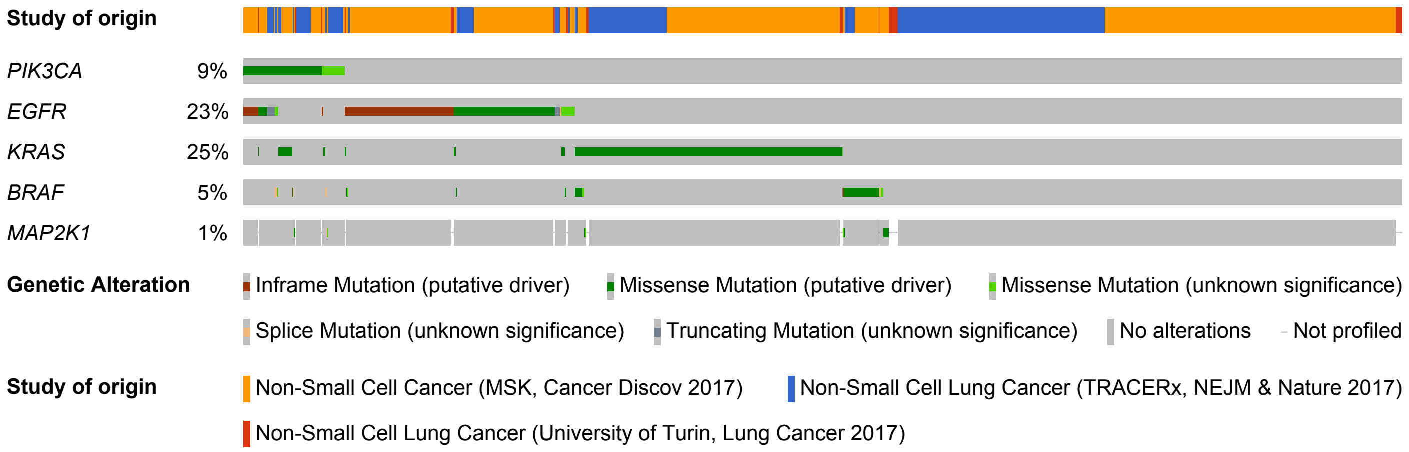


| a | b | Neither | a Not b | b Not a | Both | Log2 Odds Ratio | p-Value | q-Value | Tendency |
|---|---|---|---|---|---|---|---|---|---|
| *PIK3CA* | *EGFR* | 1002 | 79 | 278 | 44 | 1.005 | <0.001 | 0.004 | **Co-occurrence** |
| *PIK3CA* | *MAP2K1* | 1233 | 115 | 10 | 4 | 2.101 | 0.028 | 0.056 | Co-occurrence |
| *PIK3CA* | *BRAF* | 1213 | 115 | 67 | 8 | 0.333 | 0.528 | 0.538 | Co-occurrence |
| *PIK3CA* | *KRAS* | 946 | 103 | 334 | 20 | -0.863 | 0.017 | 0.055 | Mutual exclusivity |

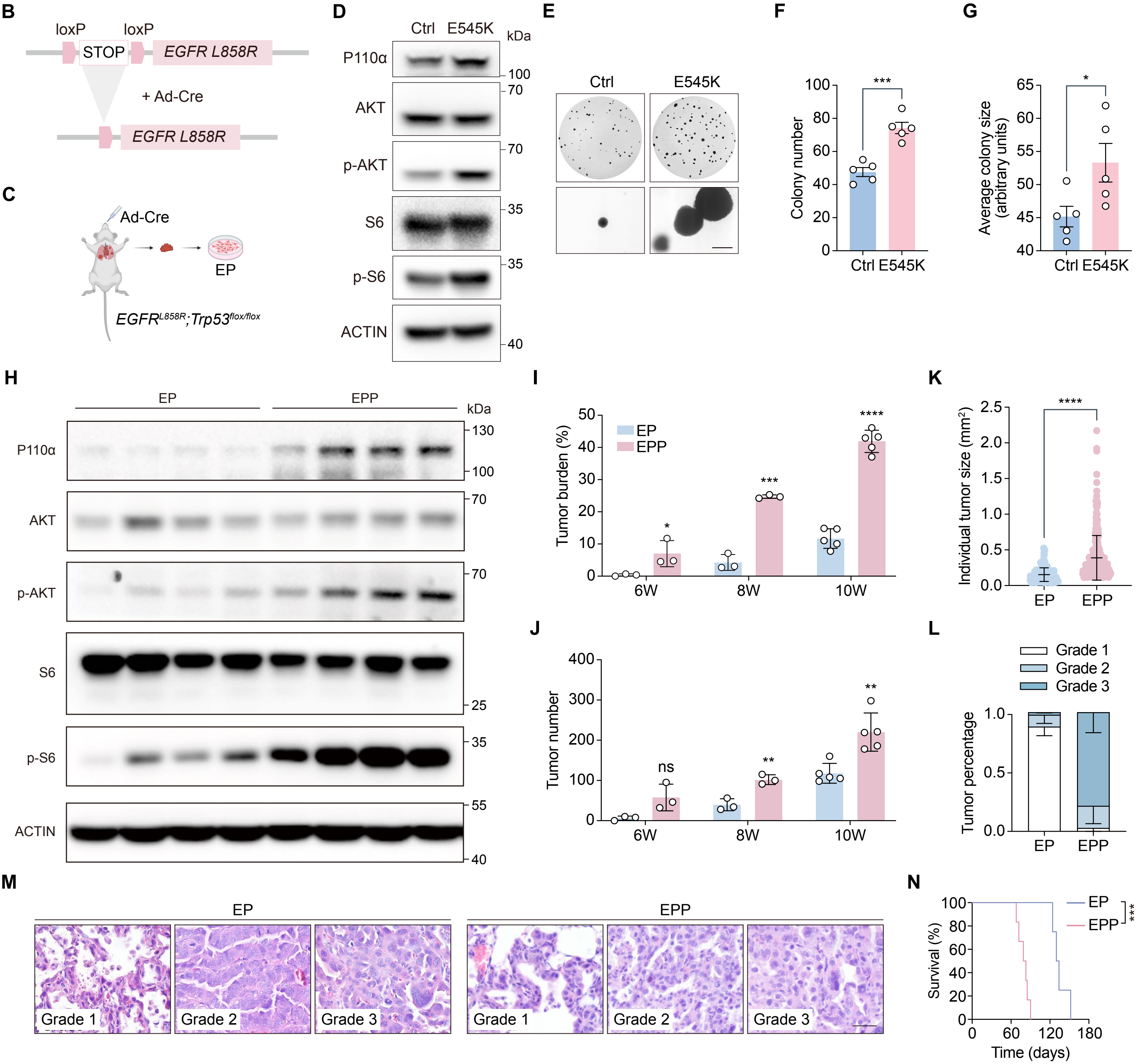

#### Figure S3

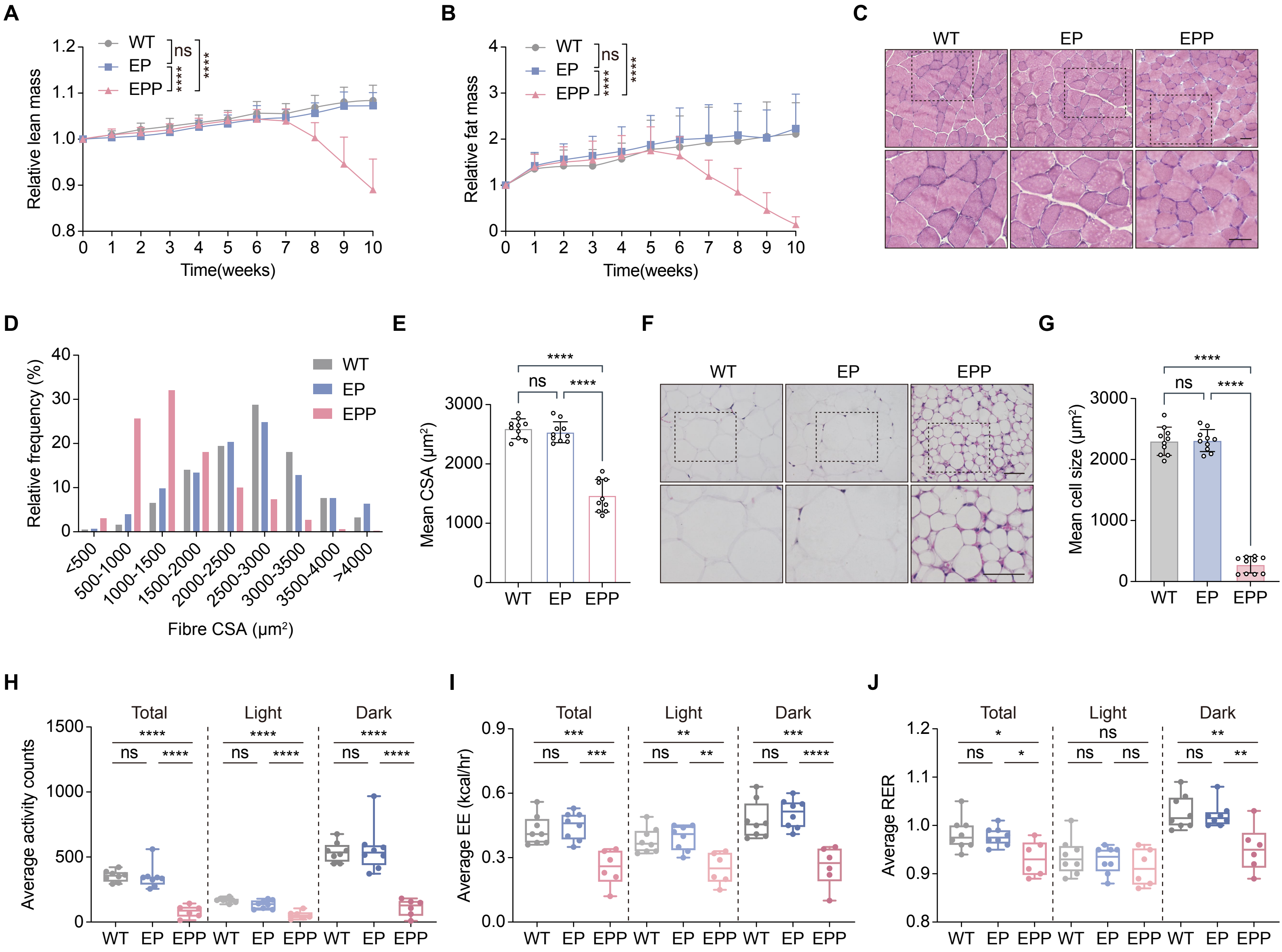
A
B
C
D
E
F
G
H
I
J
Relative lean mass
Relative fat mass
Time(weeks)
WT
EP
EPP
ns
****
Relative frequency (%)
Fibre CSA (μm²)
<500
500-1000
1000-1500
1500-2000
2000-2500
2500-3000
3000-3500
3500-4000
>4000
Mean CSA (μm²)
Mean cell size (μm²)
Total
Light
Dark
Average activity counts
Average EE (kcal/hr)
Average RER
***
**
*

**Figure S4**

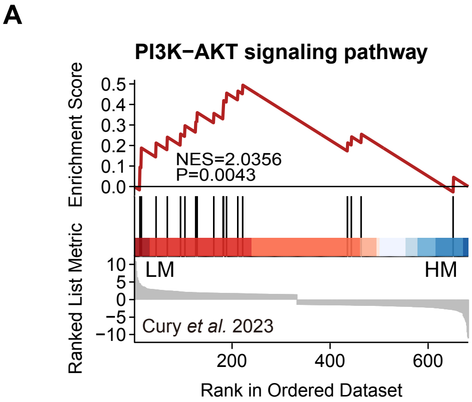
A
PI3K−AKT signaling pathway
Enrichment Score
0.5
0.4
0.3
0.2
0.1
0.0
NES=2.0356
P=0.0043
Ranked List Metric
10
5
0
−5
−10
LM
HM
Cury et al. 2023
200
400
600
Rank in Ordered Dataset

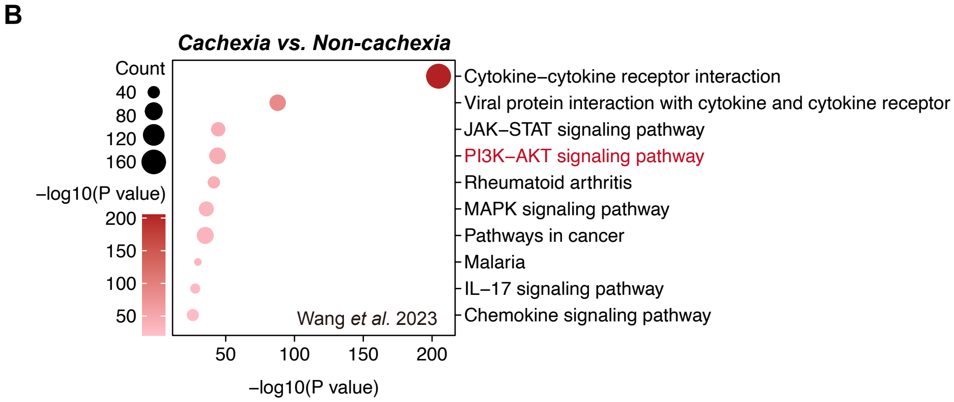
B
Cachexia vs. Non-cachexia
Count
40
80
120
160
−log10(P value)
200
150
100
50
Cytokine−cytokine receptor interaction
Viral protein interaction with cytokine and cytokine receptor
JAK−STAT signaling pathway
PI3K−AKT signaling pathway
Rheumatoid arthritis
MAPK signaling pathway
Pathways in cancer
Malaria
IL−17 signaling pathway
Chemokine signaling pathway
Wang et al. 2023
50
100
150
200
−log10(P value)

#### Figure S5

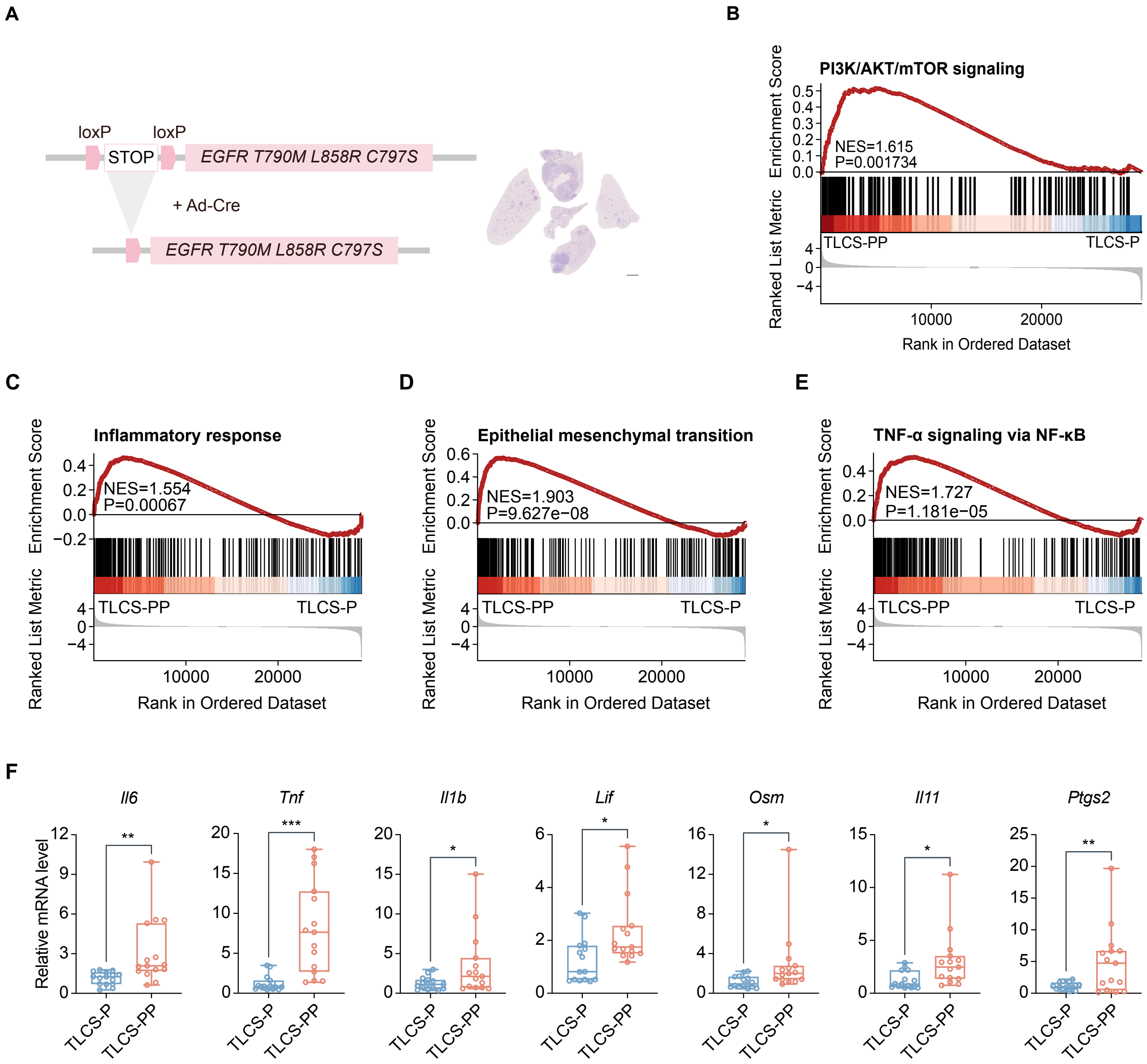

A
loxP
loxP
STOP
EGFR T790M L858R C797S
+ Ad-Cre
EGFR T790M L858R C797S
B
PI3K/AKT/mTOR signaling
NES=1.615
P=0.001734
TLCS-PP
TLCS-P
Enrichment Score
Ranked List Metric
Rank in Ordered Dataset
C
Inflammatory response
NES=1.554
P=0.00067
D
Epithelial mesenchymal transition
NES=1.903
P=9.627e−08
E
TNF-α signaling via NF-κB
NES=1.727
P=1.181e−05
F
Il6
Tnf
Il1b
Lif
Osm
Il11
Ptgs2
Relative mRNA level
TLCS-P
TLCS-PP

#### Figure S6

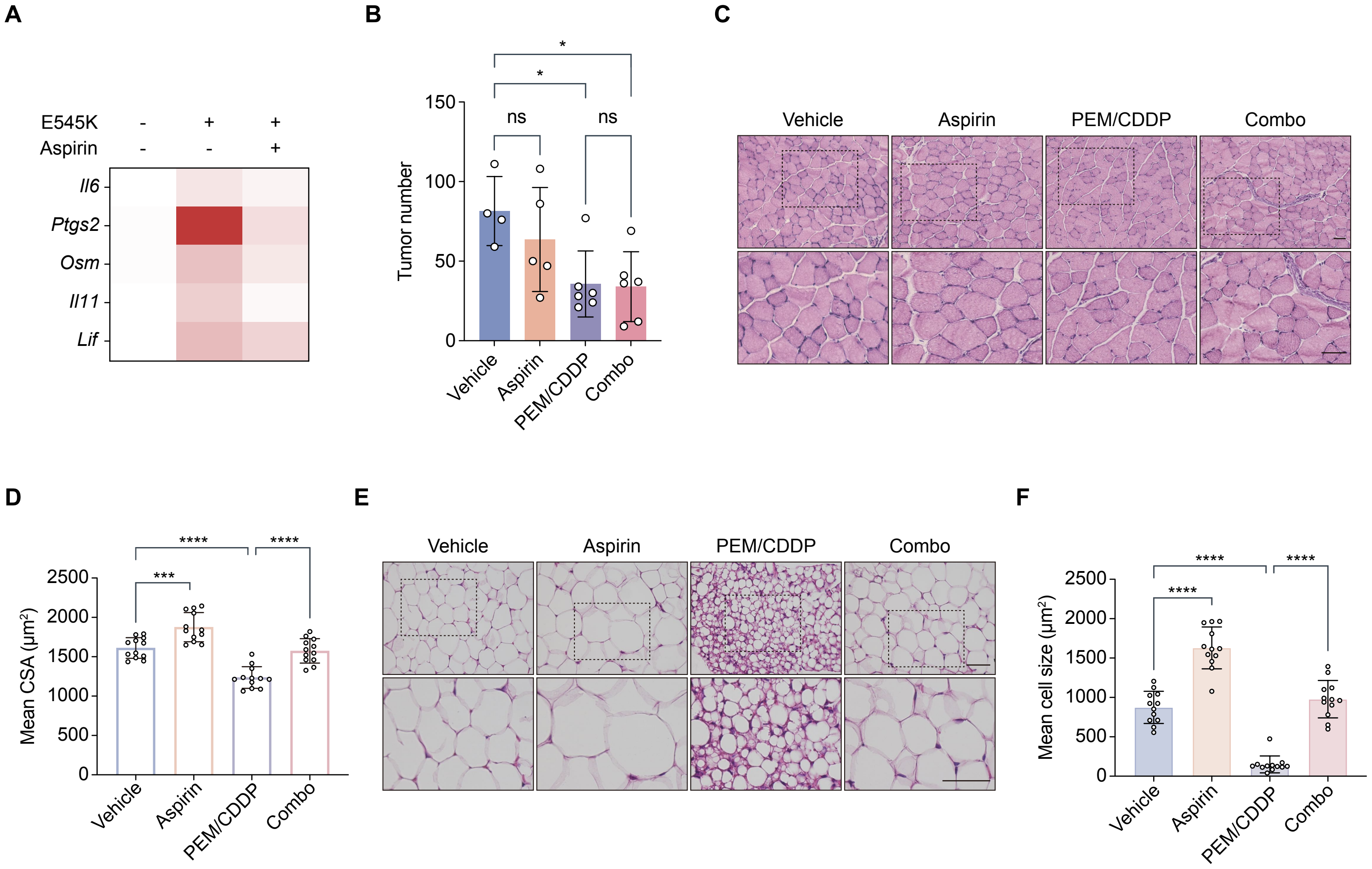

A
E545K - + +
Aspirin - - +
Il6
Ptgs2
Osm
Il11
Lif
B
Tumor number
150
100
50
0
ns
ns
*
*
Vehicle
Aspirin
PEM/CDDP
Combo
C
Vehicle
Aspirin
PEM/CDDP
Combo
D
Mean CSA (μm²)
2500
2000
1500
1000
500
0
***
****
****
Vehicle
Aspirin
PEM/CDDP
Combo
E
Vehicle
Aspirin
PEM/CDDP
Combo
F
Mean cell size (μm²)
2500
2000
1500
1000
500
0
****
****
****
Vehicle
Aspirin
PEM/CDDP
Combo
